# Lipopolysaccharide integrity primes bacterial sensitivity to a cell wall-degrading intermicrobial toxin

**DOI:** 10.1101/2023.01.20.524922

**Authors:** Kristine L Trotta, Beth M Hayes, Johannes P Schneider, Jing Wang, Horia Todor, Patrick Rockefeller Grimes, Ziyi Zhao, William L Hatleberg, Melanie R Silvis, Rachel Kim, Byoung Mo Koo, Marek Basler, Seemay Chou

## Abstract

Gram-negative bacteria can antagonize neighboring microbes using a type VI secretion system (T6SS) to deliver toxins that target different essential cellular features. Despite the conserved nature of these targets, T6SS potency can vary across recipient species. To understand the molecular basis of intrinsic T6SS susceptibility, we screened for essential *Escherichia coli* genes that affect its survival when antagonized by a cell wall-degrading T6SS toxin from *Pseudomonas aeruginosa*, Tae1. We revealed genes associated with both the cell wall and a separate layer of the cell envelope, surface lipopolysaccharide, that modulate Tae1 toxicity *in vivo*. Disruption of lipopolysaccharide synthesis provided *Escherichia coli (Eco)* with novel resistance to Tae1, despite significant cell wall degradation. These data suggest that Tae1 toxicity is determined not only by direct substrate damage, but also by indirect cell envelope homeostasis activities. We also found that Tae1-resistant *Eco* exhibited reduced cell wall synthesis and overall slowed growth, suggesting that reactive cell envelope maintenance pathways could promote, not prevent, self-lysis. Together, our study highlights the consequences of co-regulating essential pathways on recipient fitness during interbacterial competition, and how antibacterial toxins leverage cellular vulnerabilities that are both direct and indirect to their specific targets *in vivo*.

## INTRODUCTION

Many bacteria live in mixed-species microbial communities where they compete with each other for limited space and resources^1^. Intermicrobial competition is mediated by a diverse array of molecular strategies that can exclude or directly interfere with other microbes, both near and far^2^. Nearly 25% of Gram-negative bacteria encode a type VI secretion system (T6SS)^3^, which antagonizes neighboring cells by injection of toxic protein effectors into a recipient cell^4–6^. The opportunistic human pathogen *Pseudomonas aeruginosa* (*Pae*) harbors an interbacterial T6SS (H1-T6SS)^7^ that can kill the model bacterium *Escherichia coli* (*Eco*)^8–10^. Studies of H1-T6SS-mediated competition between these genetically tractable species have provided fundamental insights into the molecular basis of T6SS function and regulation.

Key to *Pae* H1-T6SS toxicity are its seven known effectors, each with a unique biochemical activity^6, 11–15^. The T6S amidase effector 1 (Tae1) from *Pae* plays a dominant role in H1-T6SS-dependent killing of *Eco* by degrading peptidoglycan (PG), a structural component of the cell wall that is critical for managing cell shape and turgor^16, 17^. Early efforts to understand Tae1 toxicity focused on its *in vitro* biochemical activity against PG, which offered key insights about how H1-T6SS targets select bacterial species. Tae1 specifically digests γ-D-glutamyl-meso-2,6-diaminopimelic acid (D-Glu-*m*DAP) peptide bonds, which are commonly found in PG from Gram-negative bacteria ^8, 18^. Tae1 toxicity is further restricted to non-kin cells through a *Pae* cognate immunity protein, T6S amidase immunity protein 1 (Tai1), which binds and inhibits Tae1 in kin cells^11, 19, 20^.

However, biochemical specificity is not sufficient to explain the toxicity and organismal selectivity of T6SS effectors *in vivo*. Bacteria antagonized by T6SSs (‘recipients’) can actively regulate effector toxicity through adaptive stress responses. *Eco* upregulates its envelope stress responses Rcs and BaeSR after exposure to the *Vibrio cholerae* (V52) T6SS effectors TseH (a PG hydrolase)^21^ and TseL (a lipase)^22^, suggesting that *Eco* could counter cell envelope damage by re-enforcing its surface^23^. Similarly, *Bacillus subtilis* triggers protective sporulation in response to a *Pseudomonas chlororaphis* (PCL1606) T6SS effector, Tse1 (a muramidase)^24^. Additional recipient-cell coordinators of T6SS effector toxicity include reactive oxygen species^25^ and glucose-dependent gene expression^26^. These studies demonstrate that T6SS effector toxicity *in vivo* may also depend on downstream adaptive features of recipient cells.

The cell wall is a complex and dynamic substrate that is actively regulated to protect the cell^27–32^, yet *Eco* is highly susceptible to lysis by Tae1 *in vivo*. We therefore hypothesized that Tae1 activity promotes H1-T6SS-mediated lysis in *Eco* through a unique strategy to overcome neutralization by the recipient cell. In this study, we investigated the *Eco* cellular features that drive its intrinsic sensitivity to H1-T6SS and the Tae1 toxin during interbacterial competition with *Pae*. Many T6SS effectors target essential cell features, so we screened the entire complement of essential *Eco* genes (plus some conditionally essential PG genes) for Tae1 susceptibility determinants. This approach complements previous genetic screens for T6SS recipient fitness which focused on nonessential gene candidates^33, 34^. While cell wall-related genes indeed impacted *Eco* susceptibility to Tae1, we also discovered a strong relationship between survival and another component of the cell envelope, lipopolysaccharide (LPS). Perturbation of LPS synthesis genes *msbA* and *lpxK* rendered *Eco* conditionally resistant to lysis by Tae1 from *Pae*. Our work revealed that LPS-related resistance was mediated through cell-biological processes that were independent of the biochemical Tae1–PG interaction. Our findings suggest that beyond biochemical specificity and adaptive stress responses lies a role for essential homeostatic processes in defining T6SS effector toxicity *in vivo*.

## RESULTS

### Adaptation of native T6SS competitions to study *Eco* susceptibility to Tae1

We developed an *in vivo* screen for genetic interactions between the cell wall-degrading H1-T6SS effector Tae1 from *Pae* and the model target bacterium *Eco*. Our screen had two fundamental design requirements: (1) the ability to distinguish between general (T6SS-dependent) and specific (Tae1-dependent) genetic interactions, and (2) the capacity to test a broad array of target cell features. We adapted an established interbacterial competition co-culture assay between H1-T6SS-active *Pae* and *Eco*, the outcome of which is sensitive to the specific contribution of Tae1^8^. In this assay *Eco* exhibits a greater fitness advantage when competed against *Pae* missing *tae1* (*Pae^Δtae^*^1^) relative to an equivalent control strain (*Pae^WT^*) (**Figure 1a**). We hypothesized that the *Pae*:*Eco* co-culture assay could be leveraged to quantitatively compare recipient cell fitness against both Tae1 (toxin-specific fitness) and the H1-T6SS (Tae1-independent fitness) in interbacterial competition.

**Figure 1:**
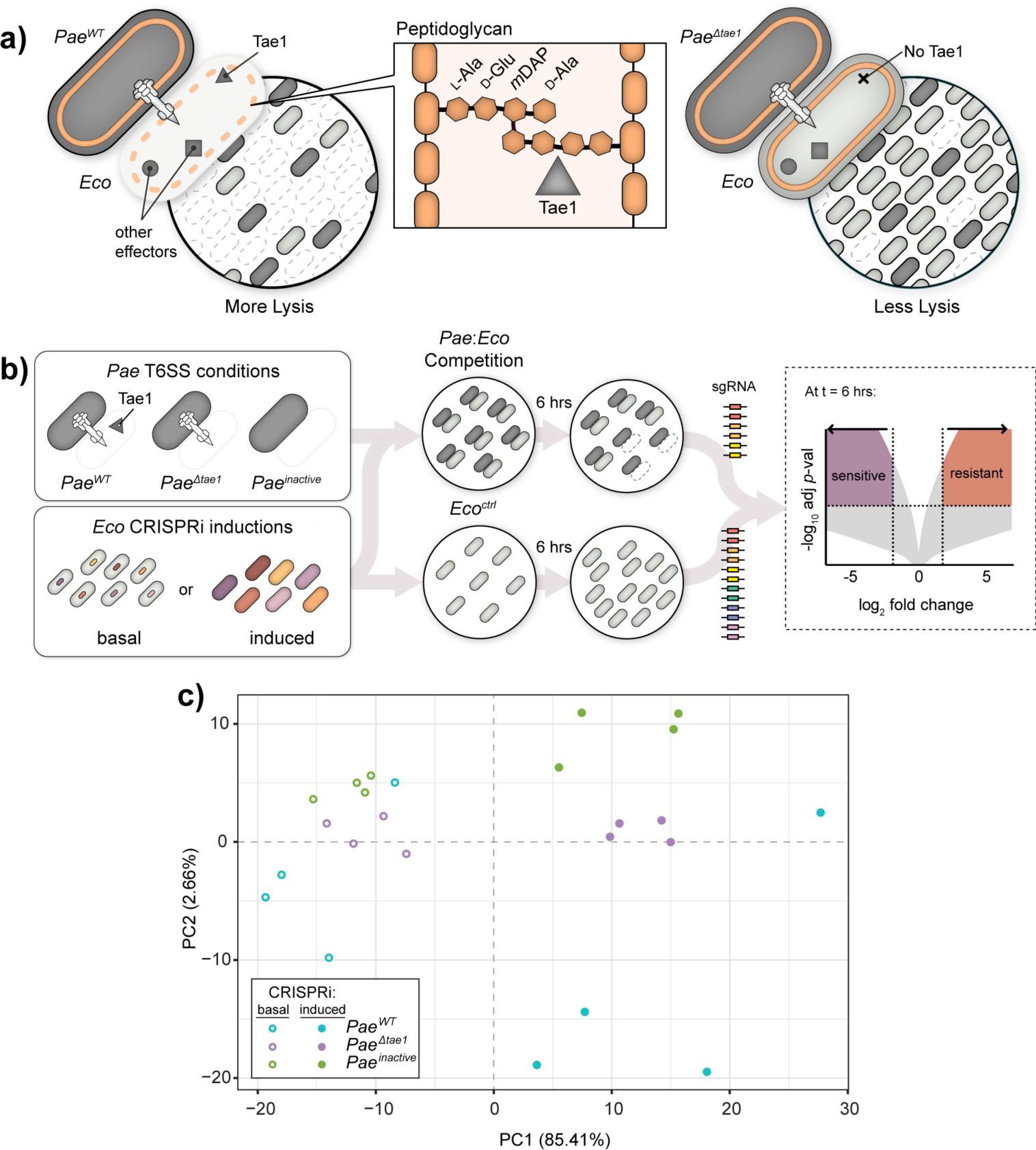
Adaptation of native T6SS competitions to study *Eco* susceptibility to Tae1. **a) Tae1 from *Pseudomonas aeruginosa* (*Pae*) degrades the *Escherichia coli* (*Eco*) cell wall to promote H1-T6SS-mediated lysis.** *Left*: *Pae^WT^* (dark grey) outcompetes *Eco* (light grey) using H1-T6SS to deliver a cocktail of toxic effectors, including Tae1 (triangle) which degrades peptidoglycan (orange). *Center*: Tae1 hydrolyzes D-Glu-*m*DAP peptide bonds in the donor stem peptides of 4,3-crosslinked peptidoglycan. *Right*: *Pae ^Δtae^*^1^ is less effective at outcompeting *Eco* using H1-T6SS. **b) Method for a genetic screen to test *Eco* gene function toward fitness against Tae1 from *Pae***. *Left*: *Pae* strains (dark grey) were engineered with modified H1-T6SS activities including: constitutively active *Pae^WT^* (*ΔretSΔpppA*), Tae1-deficient *Pae ^Δtae^*^1^(*ΔretSΔpppAΔtae1*), and T6SS-inactive *Pae^inactive^* (*ΔretSΔpppAΔicmF*). Each *Pae* strain was mixed with a pool of *Eco* KD (knockdown) strains engineered to conditionally disrupt a single gene (CRISPRi induced vs. basal). *Center*: each *Pae* strain was cocultured with an *Eco* CRISPRi strain pool for 6 hours. The *Eco* CRISPRi strain pool was also grown for 6 hours without *Pae* (*Eco^ctrl^*)as a negative control. Genomic sgRNA sequences harvested from competitions were amplified into Illumina sequencing libraries. *Right*: sgRNA barcode abundances after 6 hours were used to calculate a normalized log2 fold-change (L2FC) for each *Eco* KD strain under each condition. Above a -log10 *p*-value cutoff, a positive L2FC value indicates a KD strain which is resistant to a given condition relative to WT *Eco*; a negative L2FC value indicates a KD strain which is sensitive to a given condition relative to WT *Eco*. **c) Interbacterial competition and CRISPRi induction have distinct effects on the composition of the *Eco* CRISPRi strain library.** Principal component analysis of *Eco* library composition after competition against *Pae^WT^* (blue), *Pae^Δtae^*^1^ (purple), or *Paeinactive* (green), with induced (solid circles) or basal (hollow circles) CRISPRi induction. Four biological replicates per condition.

To screen broadly for *Eco* determinants, we adopted an established *Eco* CRISPR interference (CRISPRi) platform that generates hypomorphic mutants through intermediate gene expression knockdowns (KDs)^35^. In contrast to knock-outs or transposon mutagenesis studies, CRISPRi is systematically amenable to essential genes and thus provided an opportunity to make unique insights about genes that are typically challenging to screen for. This includes many essential (or conditionally essential) genes related to peptidoglycan (PG) metabolism, whose KDs we predicted would impact Tae1 toxicity. In this CRISPRi system, inducible sgRNA expression is coupled with constitutive dCas9 expression to conditionally repress transcription at specific loci with and without induction (“induced” and “basal” CRISPRi, respectively) (**Supplemental Figure 1a**). In total, our CRISPRi collection was composed of 596 *Eco* strains with KDs representing most cellular functions as defined by the NCBI clusters of orthologous genes (COG) system (**Supp. Fig. 1b**). Our collection also included 50 negative control strains with non-targeting sgRNAs, including *rfp-KD*, to ensure CRISPRi alone did not impact inherent *Eco* susceptibility to *Pae* (**Supp. Fig. 2a**).

**Figure 2:**
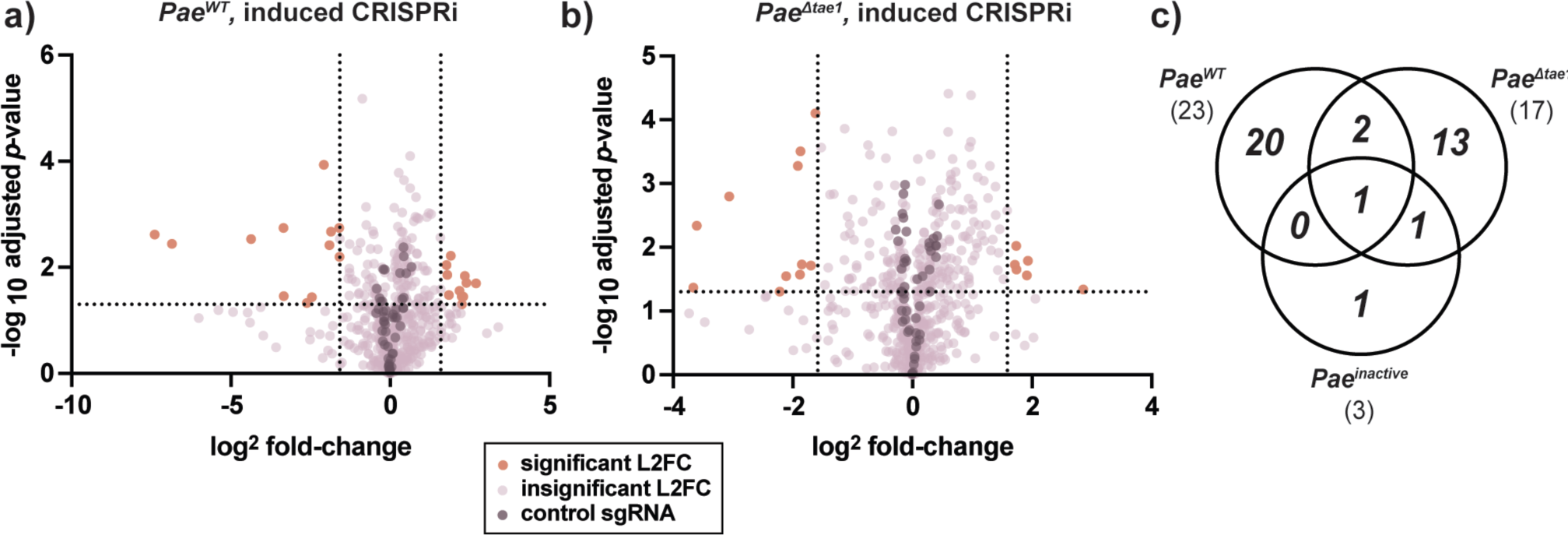
CRISPRi reveals toxin-specific and non-specific determinants of *Eco* fitness against H1-T6SS. **a-b) CRISPRi knockdowns promote *Eco* survival against *Pae^WT^*(a) and *Pae^Δtae^*^1^(b).** Volcano plots showing log2-fold change (L2FC) values for each KD strain after interbacterial competition (induced CRISPRi). Data shown: mean from four biological replicates. Statistical test: Wald test. Vertical dotted lines indicate arbitrary cutoffs for L2FC at × =-1.58 and ×=1.58 (absolute FC ×=-3 or ×= 3). Horizontal dotted line indicates statistical significance cutoff for log10 adjusted p-value (≤ 0.05). Orange points represent KDs with L2FC ≥ 1.58 or ≤ -1.58 and log10-adj. *p*-value ≤0.05. Dark purple points represent non-targeting negative control KDs (*n*=50). Lavender points represent KDs that do not meet cutoffs for L2FC or statistical test. **c) T6SS competitions identify CRISPRi strains with distinct fitness changes against T6SS and Tae1.** Venn diagram of total KDs significantly enriched OR depleted after competition against *Pae^WT^* (*n*=23), *Pae^Δtae^*^1^(*n*=17), and *Pae^inactive^*(*n*=5).

For the interbacterial competition screen, we co-cultured *Pae* with the pooled *Eco* CRISPRi collection to test competitive fitness across all KD strains in parallel (**Fig. 1b**). To compare Tae1-dependent and - independent fitness determinants, we conducted screens against H1-T6SS-active *Pae* strains that either secrete Tae1 (*Pae^WT^*; *ΔretSΔpppA*) or are Tae1-deficient (*PaeΔtae1*; *ΔretSΔpppAΔtae1*). As negative controls, we also competed the *Eco* collection against a genetically H1-T6SS-inactivated *Pae* strain (*Paeinactive*; *ΔretSΔpppAΔicmF*) and included a condition in which the collection was grown without *Pae* present (*Eco^ctrl^*). Experiments were performed under both induced and basal CRISPRi conditions to distinguish between general *Eco* fitness changes and those due to transcriptional knockdown. We used high-throughput sequencing to quantify KD strain abundance at the beginning and end of each six-hour competition. To understand the contribution of each KD to *Eco* survival against *Pae* in the presence or absence of H1-T6SS or Tae1, we calculated log2 fold-change (L2FC) values for each KD strain after competition and normalized against abundance after growth without competition (*Eco^ctrl^*)^36, 37^. Across four biological replicates per condition, L2FC values were reproducible (**Supp. Fig. 3a**; median Pearson’s *r* between all replicates = 0.91). L2FC was used as a proxy for competitive fitness of KD strains across different competition conditions.

**Figure 3:**
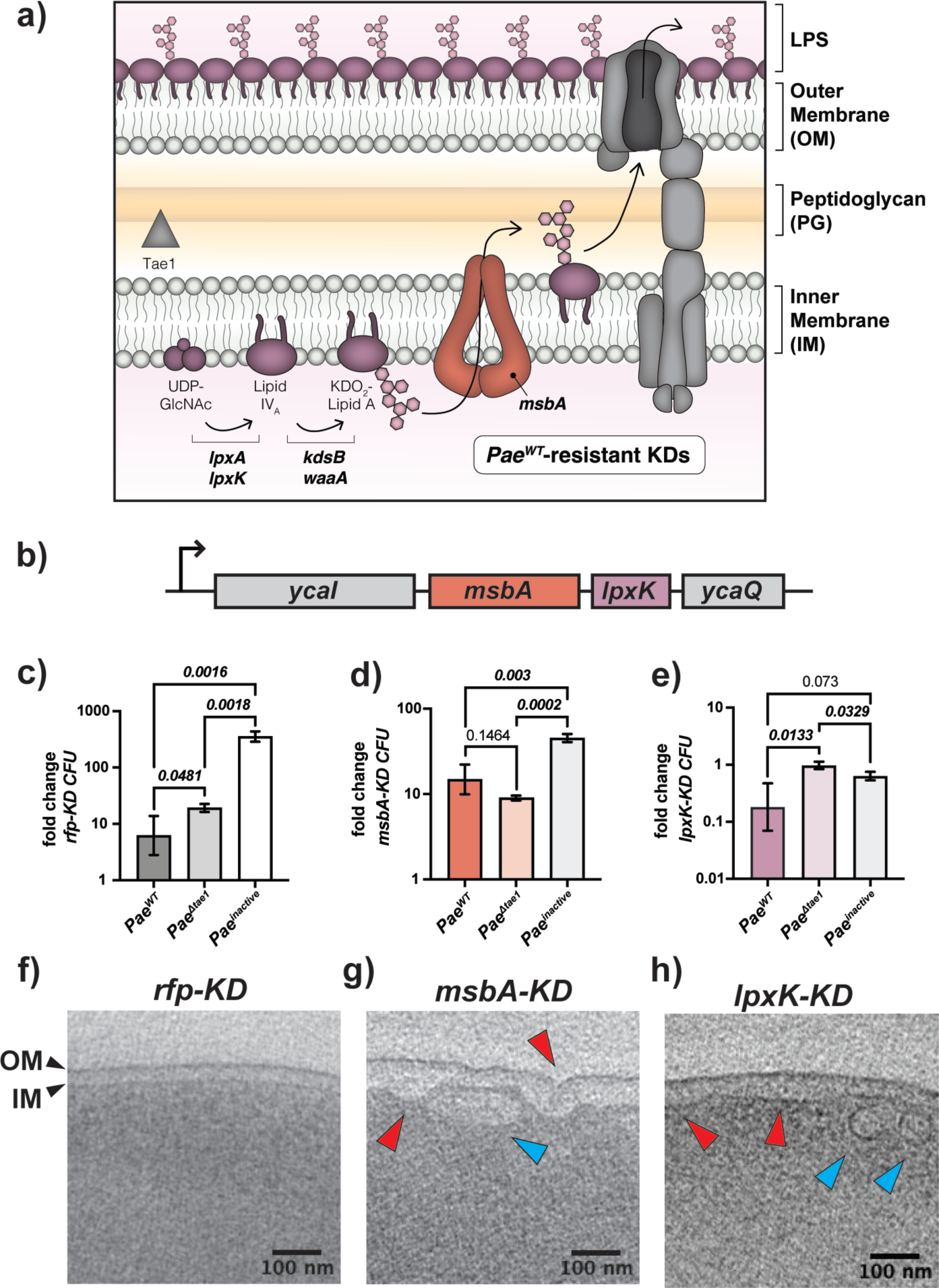
*msbA-KD* disrupts LPS biosynthesis and imparts Tae1 resistance. **a) Tae1 resistance emerges in KDs that target the lipopolysaccharide (LPS) biosynthesis pathway.** Schematic representation of the LPS biosynthesis pathway and its distribution across the *Eco* cell envelope. Genes with KDs that render *Eco* resistant to *Pae^WT^*are involved in the biosynthesis of Kdo2-Lipid A (*lpxA, lpxK, kdsA, waaA, msbA*). Note that Tae1 (grey triangle) targets peptidoglycan (PG), which is physically separate from Kdo2-Lipid A synthesis in the IM. **b) The Kdo2-Lipid A biogenesis genes *msbA* and *lpxK* are integral members of the *ycaI-msbAlpxK-ycaQ* operon in *Eco***. *msbA* (orange) and *lpxK* (purple) are co-expressed at the transcriptional level. **c-e) *msbA-KD* loses sensitivity to Tae1 in interbacterial competition against *Pae* but *lpxK-KD* does not.** Interbacterial competitions between *Pae* (*Pae^WT^, Pae^Δtae^*^1^, *Pae^inactive^*) and *rfp-KD* (c; grey), *msbA-KD* (d; orange), or *lpxK-KD* (e; purple). Data shown are average fold-change in *Eco* colony forming units (CFUs) after 6 hours of competition (geometric mean 3 biological replicates ± s.d). Statistical test: unpaired two-tailed *t-*test; *p*-value ≤0.05 displayed in bold font. **f-h) Kdo2-Lipid A mutants develop structural damage to membranes.** Cryo-EM tomographs of *rfp-KD* (f), *msbA-KD* (g), and *lpxK-KD* (h) with CRISPRi induced, highlighting cross-sections of the cell envelope (including IM and OM; black arrows). Deformed membranes (red arrows) and novel intracellular vesicles (blue arrows) are demarcated in (g) and (h). Scale bar: 100nm.

To determine if our screen was sensitive to the effects of Tae1, H1-T6SS, and CRISPRi, we conducted a principal component analysis of L2FC values for each strain under every competition condition (**Fig. 1c**). We observed clear separation of datasets by CRISPRi induction (induced versus basal) across the first principal component (PC1; 85.41%), indicating that KD induction was a major contributor to the performance of the KD library in the pooled screen. We also observed clustering of datasets according to *Pae* competitor (PC2; 2.66%). These results indicate that each *Pae* competitor yielded a distinct effect on the fitness of the CRISPRi library and demonstrates that our screen was sensitive to the presence (*Pae^WT^*) or absence (*Pae^Δtae^*^1^) of Tae1 delivery from H1-T6SS. From these data we conclude that our screen successfully captured the unique impacts of CRISPRi, Tae1, and H1-T6SS on pooled *Eco* CRISPRi libraries during interbacterial competition.

### CRISPRi reveals toxin-specific and non-specific determinants of *Eco* fitness against H1-T6SS

To reveal specific *Eco* genes that shape intrinsic susceptibility to H1-T6SS-mediated antagonism, we identified KD strains which were significantly depleted or enriched at least three-fold (L2FC<-1.585 for depletion or L2FC>1.585 for enrichment, and -log10 p-adj <0.05) after competition against *Pae^WT^, Pae^Δtae^*^1^*, or Pae^inactive^*. Our goal was to prioritize KDs which had a unique effect on fitness against *Pae^WT^* relative to conditions lacking Tae1. With CRISPRi induced, we found a select cohort of KDs with significant loss of fitness (*n*=12) or gain of fitness (*n*=11) against *Pae^WT^* (**Fig. 2a**). We were surprised that some KDs caused resistance to Tae1 despite the combined challenge of essential gene depletion and H1-T6SS antagonism.

Competition against *Pae^WT^* with basal CRISPRi diminished the pool of significant candidate KDs (**Supp. 4a**), reinforcing our observation that KD strains’ fitness changes against *Pae* are dependent on CRISPRi induction. Against *Pae^Δtae^*^1^ (CRISPRi induced), we observed seventeen KDs with significant fitness changes (**Fig. 2b**) which were also CRISPRi-dependent (**Supp. 4b**). These KDs were mostly distinct from those that affected *Eco* fitness against *Pae^WT^* (**Fig. 2c**). These results indicate that the presence or absence of Tae1 had a unique effect on the T6SS competition and thus had a distinct impact on KD fitness. Finally, we found few candidate KDs that affected fitness against *Pae^inactive^* regardless of CRISPRi induction condition (**Supp. 4a-b**), suggesting that most significant phenotypes were H1-T6SS-dependent, if not Tae1-dependent. In fact, L2FC values in *Pae^inactive^* and *Eco^ctrl^* datasets had high correlation (**Supp. 4 c-d**, median Pearson correlation *r*= 0.98), indicating that *Pae* is a neutral co-culture partner with its H1-T6SS inactivated. With our interest in Tae1-specific determinants, we focused our attention on the 20 KDs which had a unique effect on *Eco* fitness against Tae1 (*Pae^WT^* +CRISPRi induced; **Table 1**). Most KDs in this group targeted genes related to the cell envelope (COG category M: cell wall/membrane/envelope biogenesis, *n*=13/20). Composed of concentric layers of inner membrane (IM), cell wall PG, outer membrane (OM), and lipopolysaccharide (LPS)^38^(**Fig. 3b**), the cell envelope is a critical structure for protecting *Eco* against environmental stress. Tae1-sensitized strains were dominated by gene targets related to the synthesis of PG (*murA, ftsI, murC, murI, mcrB, murJ*). Given that Tae1 targets the cell wall, these results support our initial hypothesis that PG structural integrity or composition are direct determinants of Tae1 *susceptibility.* KDs related to lipid membrane metabolism and transport offered either resistance (*pssA, acpP, ffs, ffh)* or sensitivity (*accD, bamA*) to Tae1, indicating that cell envelope factors indirect to the effector-substrate interaction could impact Tae1 toxicity. To our surprise, most KDs that rendered *Eco* resistant to *Pae^WT^* were related to LPS synthesis and transport. Tae1 is not known to directly interact with the IM, OM, or LPS as part of its molecular mechanism but metabolic crosstalk does occur between the PG, LPS, and lipid biosynthesis pathways^31, 39^. Thus, our data raised the possibility that regulation of other cell envelope structures could also be implicated in mediating cell wall attack.

**Table 1:** Cell envelope gene KDs develop strong fitness changes against Tae1 in competition. KDs that target PG synthesis can increase *Pae^WT^* sensitivity, while targeting other cell envelope processes can result in sensitivity or resistance. Data shown: normalized L2FC values for all 20 KD strains with unique and significant fitness changes against *Pae^WT^* (which secretes Tae1); average of four biological replicates.

| KD target | pathway/process | Avg. L2FC (*Pae<sup>WT</sup>*) | fitness against *Pae<sup>WT</sup>* |
| --- | --- | --- | --- |
| *murA* | PG synthesis | -7.40 | sensitive |
| *ftsI* | Cell division | -6.85 | sensitive |
| *accD* | Lipid metabolism | -4.37 | sensitive |
| *lptC* | LPS transport | -3.35 | sensitive |
| *murC* | PG synthesis | -2.61 | sensitive |
| *bamA* | OM protein assembly | -2.46 | sensitive |
| *murI* | PG synthesis | -1.86 | sensitive |
| *mrcB* | PG synthesis | -1.60 | sensitive |
| *murJ* | PG transport | -1.59 | sensitive |
| *pssA* | Lipid metabolism | 1.77 | resistant |
| *hemE* | Heme metabolism | 1.79 | resistant |
| *msbA* | LPS transport | 1.84 | resistant |
| *waaA* | LPS synthesis | 1.91 | resistant |
| *lpxA* | LPS synthesis | 2.18 | resistant |
| *ffs* | Membrane trafficking/<br>secretion | 2.25 | resistant |
| *acpP* | Lipid metabolism | 2.25 | resistant |
| *ffh* | Membrane trafficking/<br>secretion | 2.30 | resistant |
| *kdsB* | LPS synthesis | 2.35 | resistant |
| *lpxK (lpxK_-1as)* | LPS synthesis | 2.39 | resistant |
| *lpxK (lpxK_32as)* | LPS synthesis | 2.69 | resistant |

### *msbA-KD* disrupts LPS biosynthesis and imparts Tae1 resistance

To investigate the hypothesis that non-PG components of the cell envelope may also shape Tae1 toxicity, we focused downstream studies on Tae1-resistant KDs related to the synthesis of LPS, an essential lipidated surface sugar that offers protection and structure to the OM^40^. Candidate KDs targeted highly-conserved, essential genes in Kdo2-Lipid A synthesis and transport (*lpxA, lpxK, kdsB, waaA, msbA*) (**Fig. 3a**). Kdo2-Lipid A synthesis is the most-upstream arm of LPS biosynthesis with rate-limiting control over the entire pathway^41, 42^. In our screen, the strongest resistance phenotypes we observed were in KDs targeting *lpxK* (*lpxK_-1as* and *lpxK_32as*) (**Table 1**). LpxK is a kinase that phosphorylates the Lipid-A intermediate tetraacyldisaccharide 1-phosphate to form Lipid IVA43,44. In *Eco, lpxK* is in an operon with *msbA* (**Fig. 3b**), which encodes the IM Kdo2-Lipid A flippase MsbA^45, 46^. A KD of *msbA* (*msbA_40as*) also conferred resistance to *Pae*^WT^ in our screen(**Table 1**).

We first experimentally validated pooled screen results by individually testing *lpxK-KD* and *msbA-KD* fitness in binary competitions against *Pae*. We regenerated and validated KD strains for *lpxK* (*lpxK_-1as*; “*lpxK-KD*”) and *msbA* (“*msbA-KD*”) for use in these experiments (**Supp. 6**). Consistent with our screen, *msbA-KD* gained Tae1-specific resistance in H1-T6SS-mediated competitions (**Fig. 3d**), exhibiting loss of sensitivity to *Pae^WT^* relative to *Pae^Δtae^*^1^. In contrast, we could not validate Tae1 resistance for *lpxK-KD* (**Fig. 3e**). Like *rfp-KD* (**Fig. 3c**), *lpxK-KD* maintains sensitivity to *Pae^WT^* relative to *Pae^Δtae^*^1^. The gene expression of *msbA* and *lpxK* are co-dependent, so we were surprised that *msbA-KD* and *lpxK-KD* did not equally reproduce Tae1 resistance. However, CRISPRi-dependent phenotypes could be controlled by factors such as transcriptional polar effects or off-target CRISPRi effects. To address their phenotypic disparities, we quantified transcriptional KD efficacy and specificity for *lpxK-KD* and *msbA-KD* with qRT-PCR. For *msbA-KD* with CRISPRi induced, we found repression of *msbA* (29-fold), l*pxK* (15-fold), and *ycaQ* (3.6-fold) expression (**Supp. Fig. 6a**). Thus, owing to downstream polar effects, our *msbA-KD* strain is a KD of both LPS candidate genes, *msbA* and *lpxK*. Conversely, *lpxK-KD* only repressed *lpxK* (71-fold) and *ycaQ* (11-fold) (**Supp. Fig. 6b**), but not *msbA*. Therefore, *msbA-KD* and *lpxK-KD* yield distinct transcriptional consequences despite targeting the same operon using CRISPRi. Next, we investigated phenotypic consequences of inducing CRISPRi in *msbA-KD* and *lpxK-KD* by comparing their cellular morphologies with cryo-electron tomography. Disruption of *msbA* and *lpxK* typically leads to structural deformation in the *Eco* cell envelope from aberrant accumulation of Kdo2-Lipid A intermediates in the IM^44, 46, 47^. Unlike *rfp-KD* negative control cells (**Fig. 3e**), *msbA-KD* cells developed irregular buckling in the IM and OM (**Fig. 3f**, red arrows). We also observed vesicular or tubular membrane structures within the cytoplasm (**Fig. 3f**, blue arrows). Such structural abnormalities are consistent with physical crowding of Kdo2-Lipid A intermediates in the IM that are relieved by vesicular internalization. On the other hand, while *lpxK-KD* had a distended IM and vesicles (**Fig. 3g**, red and blue arrows), the OM appeared smooth and regular. This phenotypic divergence points to two distinct KD effects: defects in the IM (both *msbA-KD* and *lpxK-KD*) and defects in the OM (*msbA-KD* only). Together with our transcriptional analyses, these results demonstrate that *msbA-KD* and *lpxK-KD* have unique consequences for LPS integrity and Tae1 susceptibility despite targeting the same operon. We focused the remainder of our study on the validated *msbA-KD* strain which damages the IM and OM.

### Resistance to Tae1 in *msbA-KD* is independent of cell wall hydrolysis

Identifying *msbA* and *lpxK* as potential Tae1 resistance determinants provided us a chance to study mechanisms by which LPS impacts susceptibility to cell wall damage. Such mechanisms could span several scales including: direct Tae1-PG interactions (**Fig. 4**), cellular responses to Tae1 hydrolysis (**Fig. 5**), broad physiological conditions that affect mechanical lysis (**Fig. 6**), or some combination of these. To investigate, we used an orthogonal *in vivo* assay to directly test the effect of Tae1 activity in *msbA-KD* cells in the absence of *Pae* and other co-delivered H1-T6SS toxins. We measured lysis for *rfp-KD* and *msbA-KD* upon induction of exogenous wild-type Tae1 (Tae1^WT^) expression in the cell wall-containing periplasm^8,48^and found that *msbA-KD* had increased survival against Tae1^WT^ relative to *rfp-KD* (**Fig. 4a**). *Eco* resistance was dependent on Tae1 activity, as evidenced by loss of the *msbA-KD* resistance phenotype with catalytically-attenuated Tae1^C30A^ (**Fig. 4b**) and no-enzyme (empty) (**Fig. 4c**) controls. There were no major differences in Tae1 expression levels across conditions **(Supp. Fig. 7a-b),** which ruled out the possibility that fitness was tied to toxin dose. Complementation of *msbA* by overexpression partially rescued Tae1^WT^ susceptibility in *msbA-KD* (**Supp. Fig. 8a-c,g**), while *lpxK* overexpression did not (**Supp. Fig. 8d-f,h**). Given the multigenic knockdown in *msbA-lpxK-ycaQ* in *msbA-KD*, these data suggest that *msbA* is a partial determinant of Tae1 susceptibility in the strain.

**Figure 4:**
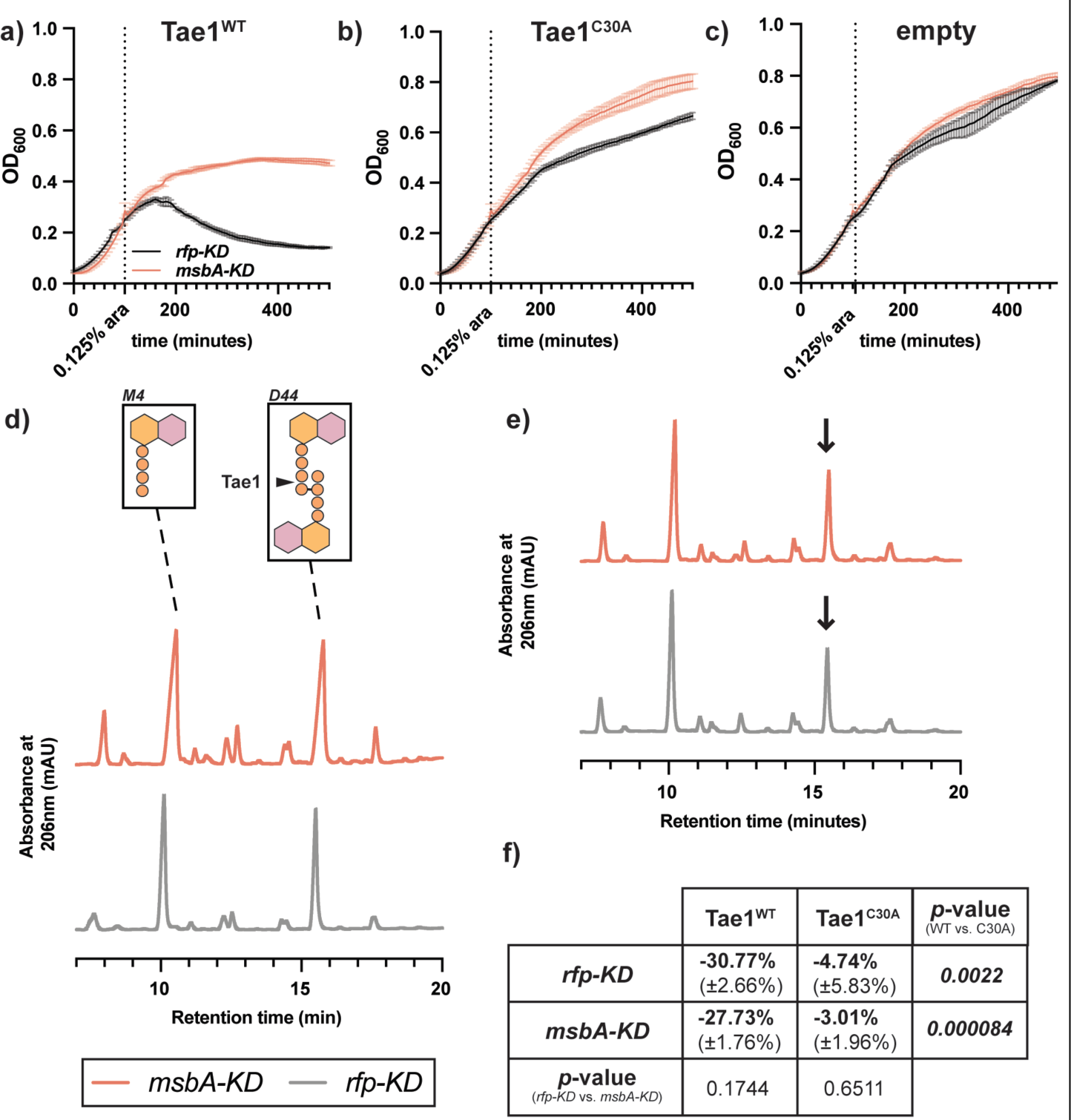
Resistance to Tae1 in *msbA-KD* is independent of cell wall hydrolysis. **a-c) *msbA-KD* populations have a Tae1-dependent growth advantage.** OD_600_ growth curves of *msbA-KD* (orange) and *rfp-KD* (black) with CRISPRi induced, overexpressing (a)*pBAD24::pelB-tae1^WT^*(Tae1^WT^), (b) *pBAD24::pelB-tae1^C30A^* (Tae1^C30A^), or (c) *pBAD24* (empty). Data shown: average of 3 biological replicates ± s.d. Dotted vertical line indicates plasmid induction timepoint (at OD_600_=0.25). **d) The muropeptide composition of *msbA-KD* PG is identical to control *rfp-KD*.** HPLC chromatograms of muropeptides purified from *msbA-KD* (orange) and *rfp-KD* (grey) expressing *pBAD24* (empty). *Inset:* major muropeptide species in *Eco* include tetrapeptide monomers (M4; r.t. ∼10 minutes) and 4,3-crosslinked tetra-tetra dimers (D44; r.t. ∼15.5 minutes). Tae1 digests D44 peptides (black arrow). Data shown: representative from 3 biological replicates. **e) Tae1^WT^ digests PG from both *msbA-KD* and *rfp-KD* PG *in vivo*.** HPLC chromatograms of muropeptides purified from *msbA-KD* (orange) and *rfp-KD* (grey) expressing *pBAD24::pelB-tae1^WT^*(Tae1^WT^). Black arrow indicates D44 peptide partially digested by Tae1. Data shown: representative from 3 biological replicates. **f) Tae1 is equally efficient at digesting PG in *msbA-KD* and *rfp-KD*.** Percent loss of D44 peptide after 60 minutes of periplasmic Tae1^WT^ or Tae1^C30A^ expression. Data shown: average of 3 biological replicates (± s.d.). Statistical test: two-tailed unpaired *t-*test; *p*-value ≤0.05 displayed in bold font.

**Figure 5:**
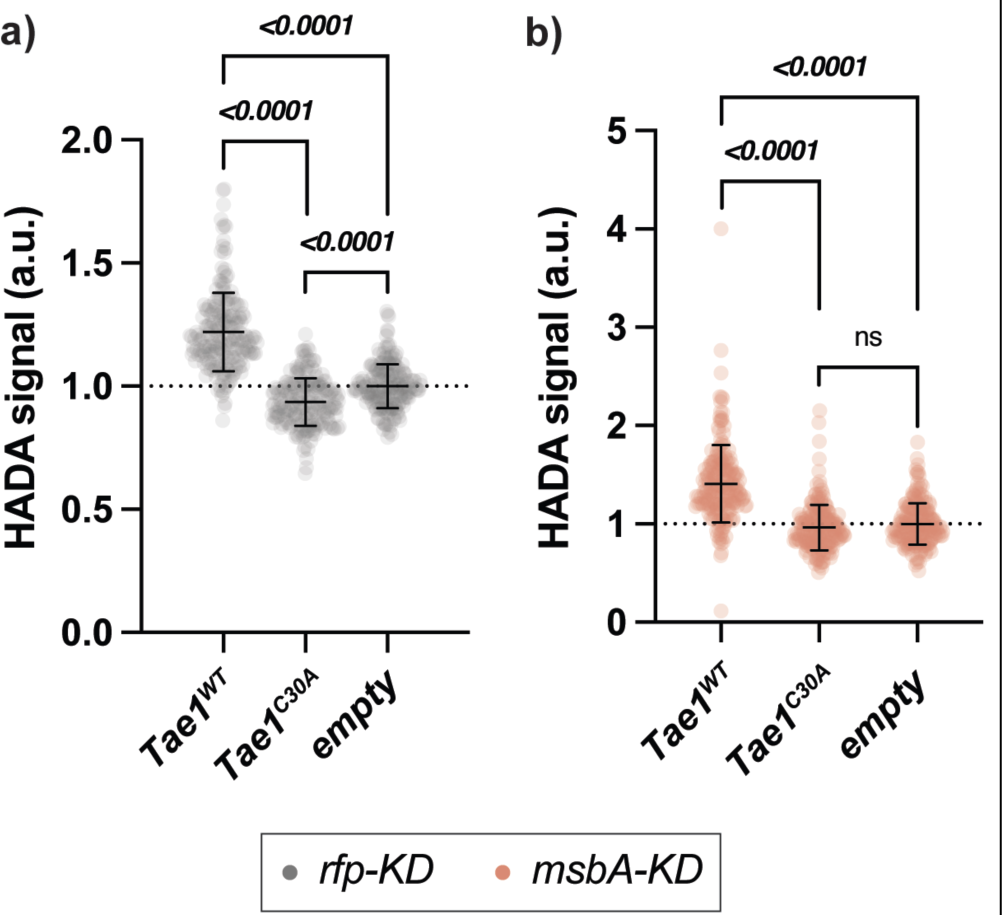
PG synthesis is suppressed in *msbA-KD* but sensitive to Tae1 activity. **a-b) PG synthesis activity is sensitive to Tae1 overexpression.** Single-cell fluorescence intensity measurements for *rfp-KD* (a; grey) or *msbA-KD* (b; orange) after incorporating the fluorescent Damino acid HADA into PG after 60 minutes of overexpressing *pBAD24::pelB-tae1^WT^* (Tae1^WT^), *pBAD24::pelB-tae1^C30A^* (Tae1^C30A^), or *pBAD24* (empty), with CRISPRi induced. Data shown: 600 cells (200 cells × 3 biological replicates), with average ± s.d. Statistical test: unpaired two-tailed *t*-test; *p*-value ≤0.05 displayed in bold font.

Next, we tested whether *msbA-KD* directly impacts Tae1–PG physical interactions by triggering changes to the chemical composition of *Eco* PG, which can occur downstream of OM stress^31^. PG remodeling could alter intrinsic Tae1 susceptibility by changing the relative abundance of targetable peptides in the cell wall. We isolated and characterized the composition of PG purified from *rfp-KD* and *msbA-KD* by HPLC muropeptide analysis. Both strains had highly similar and stereotypical *Eco* muropeptide profiles (**Fig. 4d**). PG peptides containing the scissile bond and structural context for Tae1 recognition (4,3-crosslinked dimers; D44)^8^ were found at an approximate 1:1 ratio with another dominant species of muropeptide (tetrapeptide monomers; M4)^49^. Our results suggest that the PG composition of *msbA-KD* is not modified downstream of LPS damage, indicating that Tae1 resistance cannot be explained by biochemical changes to the Tae1:PG interaction.

**Figure 6:**
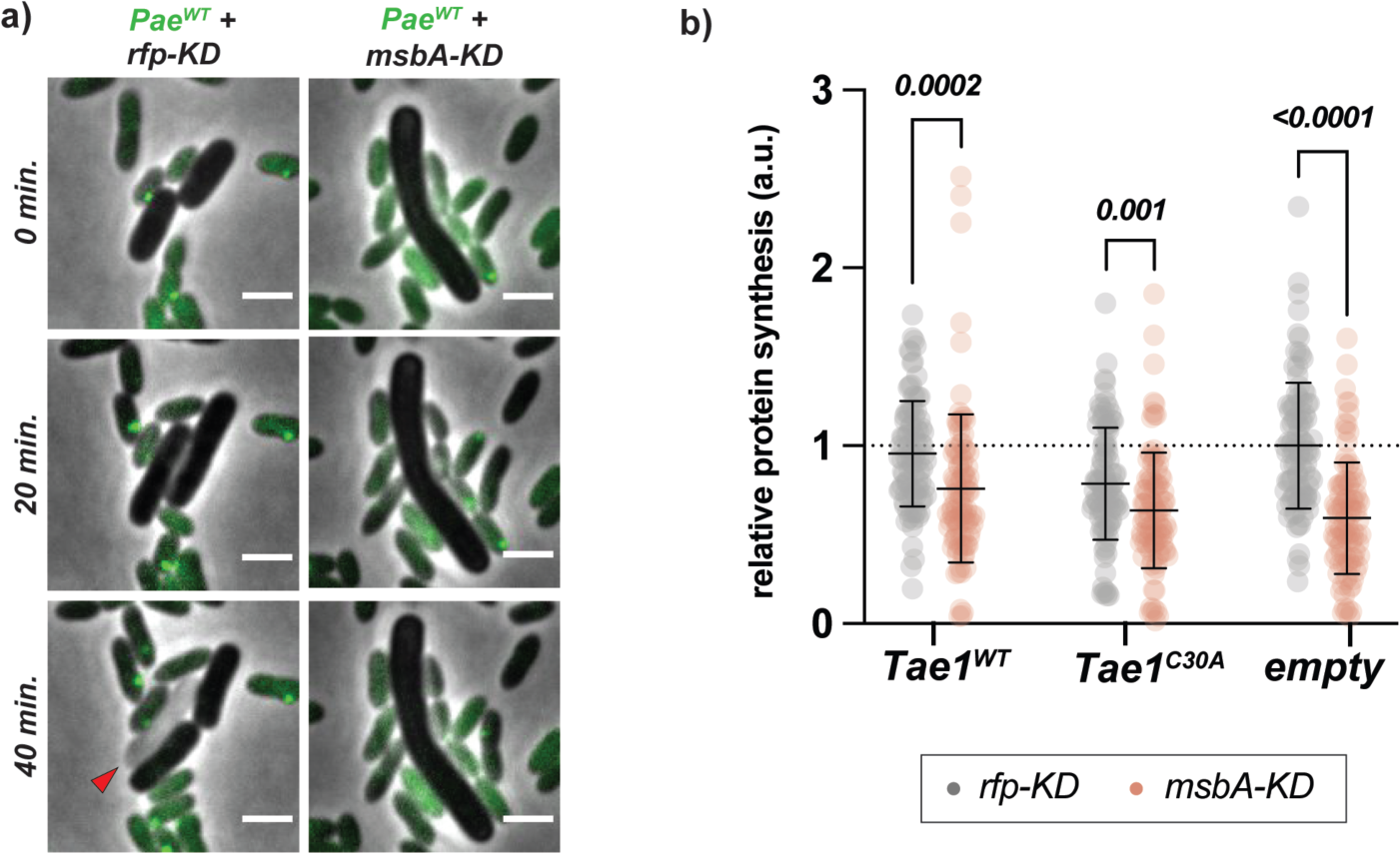
Blocks to growth and protein synthesis accompany Tae1 resistance in *msbA-KD*. **a) *msbA-KD* cells resist lysis from *Pae^WT^* while growing slowly without dividing.** Representative frames from time-course imaging of *rfp-KD* (left column; grey) and *msbA-KD* (right column; grey) co-cultured with *PaeWT* (green), with CRISPRi induced. Green foci in *Pae^WT^* indicate aggregates of GFP-labelled ClpV, which signal a H1-T6SS firing event. Red arrow indicates lysed cell. Data shown are merged phase and fluorescent channels. Scale bar: 2μm. **b) Protein synthesis activity is attenuated in *msbA-KD*.** Single-cell fluorescence intensity measurements for *rfp-KD* (grey) or *msbA-KD* (orange) cells after incorporating fluorescentlylabelled O-propargyl-puromycin (OPP) into nascent peptides during overexpression of *pBAD24::pelB-tae1^WT^*(Tae1^WT^), *pBAD24::pelB-tae1^C30A^* (Tae1^C30A^), or *pBAD24* (empty), with CRISPRi induced. All data normalized to average OPP signal in *rfp-KD +* empty. Data shown: 100 cells/condition, with average ± s.d. Statistical test: unpaired two-tailed *t*-test; *p*-value ≤0.05 displayed in bold font.

We tested an alternative hypothesis that resistance may derive from decreased efficiency in Tae1 hydrolysis. We reasoned that structural deformations in the *msbA-KD* cell envelope (**Fig. 3f**) could occlude or delay the accessibility of PG to Tae1, thus slowing the kinetics of cell wall degradation and cell lysis. To test this, we monitored the relative degradation of D44 peptides after Tae1 induction in *rfp-KD* and *msbA-KD* populations. Empty-vector and Tae1^C30A^ conditions were included as negative controls (**Fig4d,f; Supp. Fig. 9a**). At 60 minutes of induction (just prior to lysis in *rfp-KD* populations), we found that D44 peptides were similarly hydrolyzed between strains, with a 32.58% loss in *rfp-KD* and 27.73% of in *msbA-KD* (**Fig. 4e-f**). Thus, Tae1 hydrolyzes *msbA-KD* PG as efficiently as *rfp-KD* PG. Collectively, these data show that both cell wall recognition and hydrolysis by Tae1 are unchanged in *msbA-KD*, ruling out the possibility that direct changes to PG are responsible for differential cellular lysis outcomes.

### PG synthesis is suppressed in *msbA-KD* but sensitive to Tae1 activity

Given that we did not find any effects on direct Tae1–cell wall interactions in *msbA-KD*, we next explored indirect resistance mechanisms. The PG sacculus is dynamically synthesized, edited, and recycled *in vivo* to maintain mechanical support to the cell during growth and stress^27, 50^. We hypothesized that Tae1 hydrolysis could also impact PG synthesis activity in *Eco* by generating a need to replace damaged PG with new substrate. The ability to repair PG could thus be a valuable determinant of Tae1 susceptibility. To determine if PG synthesis is sensitive to Tae1 exposure, we measured the incorporation of the fluorescent D-amino acid HADA into *rfp-KD* cell walls both with and without exogenous Tae1 expression. When normalized against control cells (*empty*), PG synthesis in *rfp-KD* cells increased by 22% in response to Tae1^WT^ and decreased by 6.5% in response to Tae1^C30A^ (**Fig. 5a; Table 2**). These data show that PG synthesis is stimulated by Tae1 exposure, and this response is dependent on toxin activity.

**Table 2:** PG synthesis activity is sensitive to CRISPRi and Tae1 overexpression. Descriptive statistics for normalized percent change in HADA fluorescence in *rfp-KD* and *msbA-KD* as related to **Fig.5** and **Supp. Fig. 10**. Data shown: average of 600 single-cell measurements ±s.d.

|  |  | % change<br>(intra-strain) | % change<br>(*rfp-KD* norm.) |
| --- | --- | --- | --- |
| *rfp-KD* | Tae1<sup>WT</sup> | 22% ($\pm 3.6\%$) | |
| | Tae1<sup>C30A</sup> | -6.5% ($\pm 2.6\%$) | |
| | empty | 0% ($\pm 1.6\%$) | |
| *msbA-KD* | Tae1<sup>WT</sup> | 26.5% ($\pm 2.5\%$) | 12% ($\pm 2.5\%$) |
| | Tae1<sup>C30A</sup> | 2.82% ($\pm 3.2\%$) | -9% ($\pm 3.2\%$) |
| | empty | 0% ($\pm 2.0\%$) | -11.5% ($\pm 2.0\%$) |

PG synthesis is also coordinated to other essential processes in *Eco*, and sensitive to their genetic or chemical perturbations^31, 51^. We investigated if *msbA-KD* impacts the dynamic PG synthesis response to Tae1. Tae1^WT^ exposure yielded a 26.5% increase in PG activity in *msbA-KD*, and no significant change in activity with Tae1^C30A^ (**Fig. 5b; Table 2**). These results indicate that PG synthesis is still actively regulated in *msbA-KD* in accordance with relative Tae1 activity. However, when normalized against baseline *rfp-KD* activity, all PG synthesis measurements for *msbA-KD* were significantly diminished (**Supp. Fig. 10; Table 2**). This observation indicates that PG synthesis activity is globally suppressed as a consequence of CRISPRi in *msbA*. Thus, we conclude that PG dynamism in *Eco* is sensitive to Tae1 hydrolysis of PG, and that *msbA-KD* alters the global capacity for PG synthesis activity without altering its sensitivity to Tae1. Furthermore, these data suggest a reactive crosstalk between LPS and PG synthesis activities *in vivo*.

### Blocks to growth and protein synthesis accompany Tae1 resistance in *msbA-KD*

Based on its responsiveness to Tae1 exposure, we might hypothesize that *Eco* stimulates PG synthesis to attempt protection against lysis by Tae1. However, suppressed PG synthesis activity alongside tolerance to wildtype-levels of PG damage in *msbA-KD* suggested that *msbA-KD* may survive lysis by Tae1 using an additional strategy to support or even supersede PG integrity. *Eco* can resist lysis upon acute PG stress by transiently arresting homeostatic functions like cell division, DNA replication, and protein synthesis to prioritize stress responses to critical damage^52–54^. A recent study showed that a CRISPRi KD in *lpxA*, the first enzyme in Lipid A biosynthesis, triggered hallmark signs of a dormancy stress response called the stringent response^55^. Thus, we hypothesized that decreased PG synthesis activity in *msbA-KD* may be symptomatic of a general, KD-dependent slow growth phenotype which could protect against Tae1 activity by passive tolerance.

To observe the effects of Tae1 and CRISPRi on cellular growth and lysis behaviors over time, we performed timelapse microscopy of *rfp-KD* and *msbA-KD* cells in competition with *Pae*. Across all *Pae* competitions, *msbA-KD* cells grew slowly without dividing or lysing (**Fig. 6a; Supp. Fig. 11a-b**). By contrast, *rfp-KD* cells grew and divided rapidly, but lysed when in competition against *Pae* strains with active H1-T6SSs (*Pae^WT^, Pae^Δtae^*^1^) (**Fig. 6a; Supp. Fig. 11a-b**). These data demonstrate that stunted cell growth and division are additional consequences of CRISPRi in *msbA-KD*. We orthogonally tested the effect of *msbA-KD* on global cell physiology by measuring nascent protein synthesis activity in *msbA-KD* and *rfp-KD*. Overall protein synthesis levels were significantly lower in *msbA-KD* relative to *rfp-KD* under all conditions tested (**Fig. 6a**). From these data we conclude that *msbA-KD* cells exhibit broad changes in cellular physiology that may underscore their unique ability to survive PG damage by Tae1.

We propose a model in which Tae1 susceptibility *in vivo* is determined at multiple levels of specificity in *Eco*: not only at the level of local PG damage but also by crosstalk between essential cell envelope pathways and the general growth state of the cell. As mediated through damage to LPS in *msbA-KD*, we posit that such crosstalk between essential cell functions can be helpful for slowing reactivity and thus increasing tolerance to acute PG stress. By the same token, the enmeshment of essential pathways may render fast-growing *Eco* vulnerable to Tae1 by creating a sudden chain-reaction of imbalances in critical functions which the cell must also resolve alongside the initial PG damage.

## DISCUSSION

The species composition of mixed-microbial communities can be driven by competitive strategies that bacteria use to antagonize their neighbors. However, our understanding of microbial weapons is primarily derived from *in vitro* studies of their molecular mechanisms. In this study, we wanted to understand how Tae1, a PG-degrading H1-T6SS effector toxin, specifically aided *Pae* in antagonizing *Eco in vivo*. By combining T6SS-mediated competition with CRISPRi against essential *Eco* genes, our high-throughput genetic screen was poised to uncover new molecular details about the interaction between Tae1 and essential functions in recipient cells. Related studies have successfully identified roles for nonessential genes that contribute to recipient survival against individual T6SS effectors^33, 34^. Our study expands our understanding of intrinsic fitness against T6SS effectors by demonstrating how essential, homeostatic cell activities can have both direct (PG) and indirect (LPS, growth) impact on the effector-substrate interaction *in vivo*. We find that Tae1 toxicity is driven not only by its ability to destroy PG but also by broader physiological and regulatory contexts.

Through the lens of LPS perturbation (*msbA-KD*), we discovered that slowing cell growth is associated with resistance to Tae1-dependent lysis. The protective nature of abject dormancy has been demonstrated for survival against other cell wall-degrading enzymes, lytic bacteriophages, and antibiotics^56–60^. However, previous work on interbacterial competition has shown that fast growth protects recipient cells from T6SS by establishing stable microcolonies more quickly than T6SS can kill the recipient cell type^61, 62^. Our study suggests that slow recipient growth could also offer a fitness advantage against lytic T6SS effectors. Similarly to how dead (unlysed) cells can physically block T6SS-wielding competitors from progressing in space ^63^, slow-growing cells could also absorb T6SS attacks to protect their kin in community settings. A compelling direction for future work could be to determine if slowing cell growth by an orthogonal mechanism, such as a bonafide stringent response, is sufficient to recapitulate resistance to Tae1 or other lytic T6SS effectors.

A surprising feature of lysis resistance in *msbA-KD* was its tolerance to PG damage by Tae1 alongside additional damage to its IM and OM. Structural destabilization of the cell envelope commonly renders *Eco* hypersensitive to lysis^64, 65^. However, our observations suggest that integrity of individual envelope components is not always sufficient to explain cell lysis. Indeed, PG and the OM can work together to bear cellular turgor pressure changes by sharing the mechanical load across both surfaces^66^. The damaged OM observed in *msbA-KD* could therefore maintain its turgor-bearing properties to protect cells against lysis when Tae1 hydrolyzes PG. Additionally, the mechanical integrity of the cell envelope in *msbA-KD* may be fortified by covalently-bound Braun’s lipoprotein or changes to membrane composition which could increase cell envelope stiffness^67, 68^. Another unique feature for *msbA-KD* is that its LPS damage does not stimulate PG remodeling, unlike other depletion alleles for LPS biosynthesis affecting transport to the OM^31^. We suggest that this indicates multiple nodes for co-regulation between PG synthesis and LPS synthesis pathways with distinct phenotypic consequences. In line with this hypothesis, our screen revealed opposing Tae1 sensitivity phenotypes for KDs of *lptC* (LPS transport to OM; sensitive) and every other LPS hit from the screen (Lipid A-Kdo2 synthesis/transport; resistant). This observation invites deeper investigation into the potential for multiple types of LPS and PG crosstalk which may inform the complex underpinnings of mechanical integrity within the bacterial cell envelope.

Another key insight from our study is that PG synthesis is stimulated in response to Tae1, indicative of an active *Eco* counterresponse. However, wild-type levels of PG synthesis were coincident with, not counter to, lytic death. Diminished PG synthesis activity in *msbA-KD* could therefore enable resistance by suppressing a toxic dysregulation of homeostatic activities. We propose that Tae1 activity leads to *Eco* cell death, in part, by triggering a futile cycle of Tae1 hydrolysis and PG synthesis that does not resolve in cell wall homeostasis. An exciting prospect for future studies could involve determining the molecular mechanisms that control PG synthesis stimulation after Tae1 hydrolysis, including whether Tae1 may also synergize or hijack specific endogenous cell wall enzymes to amplify its damage to PG^69^. The dynamic regulation of PG features indirect to Tae1’s peptide target, such as the glycan backbone, interpeptide crosslinks (type and amount), and recycling could also intersect with the toxin’s acute function to affect its overall impact on the cell wall.

In conclusion, our work highlights how recipient susceptibility in interbacterial competition may be more complex than direct -substrate interactions alone. Toxins with essential targets not only impact specific molecules but also a dynamic network of interconnected pathways. T6SSs often encode multiple toxins that antagonize different essential features^70^, including components of the cell envelope and other metabolic pathways. We posit that T6SSs deploy a cocktail of toxins that can act in coordination to disrupt the network beyond repair, or even weaponize protective homeostatic mechanisms themselves. This study points to the importance of studying the role of essential genes in the context of T6SS-mediated bacterial antagonism.

## METHODS

### Bacterial growth and selection

*Escherichia coli* strains were cultured in LB or LB-no salt (LBNS) at 37°C with orbital shaking. *Pseudomonas aeruginosa* strains were cultured in LB+ 0.01% Triton at 37°C with orbital shaking. Interbacterial competitions between *Eco* and *Pae*, and all *Eco* assays requiring solid growth, were conducted on LB+agar or LBNS+agar plates at 30°C. Where necessary, bacterial strains and plasmids were selected for growth using the following antibiotics: carbenicillin (Carb; 50 µg/ml) (Grainger), chloramphenicol (Chl; 25 µg/ml) (MP Biomedicals), gentamicin (Gent; 50 µg/ml)(Alfa Aesar), irgasan (Irg; 25 µg/ml) (Sigma-Aldrich), trimethoprim (Trm;15 µg/ml) (Sigma-Aldrich), or kanamycin (Kan; 50 µg/ml.) (VWR).

### *Eco* CRISPRi library construction and use

The *Eco* CRISPRi collection was received in pooled format as a gift from the laboratory of Carol Gross (UCSF). CRISPRi strains were derived from K12 strain BW25113^71^ and are each engineered with a chromosomal insertion of *dcas9* (constitutive expression) and a custom sgRNA sequence for inducible dCas9-mediated knockdown of a single gene-of-interest^35^. Transcriptional knockdown is induced with addition of 100µM IPTG (“induced”) into growth media, though growth without inductant also results in a mild knockdown phenotype (“basal”)^35^. Except where indicated, CRISPRi knockdown is induced in this study. CRISPRi strains *msbA-KD* and *lpxK-KD* were reconstructed from the parent strain for individual use in this study. Reconstructed strains were validated by Sanger sequencing (of the sgRNA and dCas9 chromosomal inserts), qRT-PCR (for knockdown efficiency), and Western blot (for dCas9 expression). See **Table S1** for strain descriptions and **Table S2** for primer sequences used for construction and validation.

### *Pae* strain construction

*Pae^Δtae^*^1^ (*ΔretSΔpppAΔtae1; clpV-GFP*) and *Pae^inactive^* (*ΔretSΔpppAΔicmF; clpV-GFP*) strains were constructed from biparental mating of parent strain *Pae^WT^ (*B515: PAO1 *ΔretSΔpppA; clpV-GFP)*^72^ with *Eco* SM10 λpir^73^ bearing suicide vector pEXG2 cloned with homology to the gene(s) of interest and a spacer sequence for replacement. pEXG2 plasmids were cloned using splice-overlap extension^11^. After mating, transformants were isolated by negative selection on LB-agar + 5% sucrose and confirmed as scarless knockout mutants by colony PCR of the locus of interest. See **Table S1** for strain descriptions and **Table S2** for primer sequences used for construction and validation.

### Pooled interbacterial competition screen

Competition assays were performed with overnight *Pae* cultures (*Pae^WT^, Pae^Δtae^*^1^, *Pae^inactive^*) and pooled *Eco* CRISPRi libraries. Flash-frozen glycerol stocks of *Eco* pools were resuspended in LB, backdiluted to OD600=0.25, and recovered for 90 minutes at 37°C with shaking. All cultures were washed twice with fresh LB, then OD600-adjusted to 2.0 (for *Pae*) or 1.0 (for *Eco*) in either LB (basal CRISPRi) or LB+100µM IPTG (induced CRISPRi) . An aliquot of each CRISPRi pool was reserved by pelleting and flash-freezing for sequencing-based analysis of strain abundances in the starting population. Media-matched *Pae* and *Eco* were mixed at a 1:1 volumetric ratio, except for *Eco^ctrl^* populations (for which *Eco* pools were not mixed with *Pae)*. Six, 10µl aliquots of coculture were applied to nitrocellulose membranes (0.2µm, GVS) atop LB-agar (basal CRISPRi) or LB-agar +100µM IPTG (induced CRISPRi) plates to match liquid media conditions. Covering the agar surface with nitrocellulose allows for nutrient transfer from the media to the bacteria, while aiding in bacterial recovery from the surface after competition. Cocultures were dried down to the membrane under flame-sterilization, then incubated at 30°C for 6h. Cocultures were removed from the plate by scalpel-excision of surrounding nitrocellulose and resuspended into 1ml fresh PBS by bead-beating for 45s on a tabletop vortex. The six aliquots per experiment were pooled, centrifuged (2min at 9000xG, RT), and PBS was decanted. Pellets were flash frozen in liquid nitrogen and stored at -80°C.

### Sequencing library preparation

Genomic DNA was extracted from frozen bacterial pellets by phenol: chloroform extraction and RNase treatment^74^, followed by quantification on a Nanodrop 2000 spectrophotometer (Thermo Scientific). PCR amplification was used to isolate *Eco* sgRNA sequences from mixed genomic DNA and to attach Illumina Truseq index adapters for high-throughput sequencing. Sequencing libraries were purified by gel electrophoresis on 8% TBE gels (Invitrogen Novex), stained with SYBR Gold (Invitrogen) to visualize library bands, and scalpel-excised (200-300bp region) under blue light imaging (Azure Biosystems c600). Excised libraries were gel-extracted and precipitated^75^, then resuspended in nuclease-free distilled water (Invitrogen UltraPure). Library concentration was quantified on a Qubit 2.0 fluorimeter (Invitrogen) using the dsDNA high-sensitivity assay, and assayed for purity on a 2100 Bioanalyzer (Agilent) using the high-sensitivity DNA assay. Single-end sequencing was performed on an Illumina NextSeq 500 using a custom sequencing primer and a read length of 75bp. Multiplexed samples were spiked with 5% PhiX Control v3 DNA (Illumina) to account for low diversity among sgRNA sequences. See **Table S2** for custom primers used for library preparation and sequencing.

### Sequencing data analysis

Raw FASTQ files were aligned to the library oligos and counted using ScreenProcessing (https://github.com/mhorlbeck/ScreenProcessing). Counts were normalized to a total of 20,000,000 reads, pseudocounts of 1 were added, and log2 fold change (L2FC) from t0 was calculated for each strain with at least 100 counts at t0. L2FC was further corrected by subtracting the median L2FC of the non-targeting control sgRNAs from that sample^76^. The L2FC of each sgRNA were averaged across four biological replicates to calculate the L2FC for that condition. Finally, to account for differences in the number of generations experienced (growth) in each of the experimental conditions, L2FC values for the *Pae^WT^, Pae^Δtae^*^1^, *Pae^inactive^*experiments were corrected by the coefficient of a robust (MM-type) intercept free linear regression between the experimental L2FC values and the CRISPRi induction-matched (induced/basal) *Eco^ctrl^*experiment. See Table S3 for correction coefficients and corrected L2FC values. Differences between conditions were then calculated for each sgRNA as:

Diff = (L2FC [*condition*]) – (L2FC *Eco^ctrl^*)

Final Diff values are listed in **Table S4** and were used for all further analyses.

### COG analysis

Gene ontology information was compiled from the NIH Database of Clusters of Orthologous Genes (COGs) (https://www.ncbi.nlm.nih.gov/research/cog) and reported previously^35^.

### Data availability and software

Illumina sequencing data from this study is accessible at the NCBI Sequence Read Archive under accession PRJNA917770. Principal component analysis was performed using R^77^ and visualized using ggplot2^78^. All other data visualizations were prepared using GraphPad Prism 9.4.1 (GraphPad Software, San Diego, California USA, www.graphpad.com).

### Pairwise Interbacterial T6SS competition assay

Competition assays were performed with overnight liquid cultures of *Pae* and *Eco* CRISPRi strains. *Eco* cultures were backdiluted 1:4 in LB-no salt (LBNS; cite) + 100µM IPTG and grown for 1h at 37°C with shaking to pre-induce CRISPRi before competition. Strains were washed and mixed in a 1:1 volumetric ratio of *Pae* (OD600=2) and *Eco* (OD600=1) in LBNS+100µM IPTG. Three, 10µl aliquots of each liquid co-culture applied to nitrocellulose membranes (0.2µm, GVS) atop LB-agar+100µM IPTG and dried down by flame-sterilization to encourage interbacterial competition. Cocultures were incubated at 30°C for 6h. For initial *Eco* colony-forming unit measurements (CFUt=0h), 20µl of each liquid co-culture input was serially diluted (10-fold dilutions × 8) in a 96-well plate (Corning) and plated onto LB-agar + Gent (*Eco*-selective). After the competition, coculture spots were harvested from the plate by scalpel-excision of the surrounding nitrocellulose, and pooled by resuspension into 1ml fresh PBS by bead-beating for 45s on a tabletop vortex. Resuspensions were serially diluted (10×8) and plated onto LB+Gent. All serial dilution plates were incubated overnight at 37°C. Dilution plates with approximately 20-200 colonies-per-plate were counted for *Eco* CFU abundance (CFUt=0h, CFUt=6h). Fold-change in *Eco* CFUs was determined by back-calculating CFUs per ml from dilution plates, and then calculating CFUt=6h/CFUt=0h. Experiment was performed for three biological replicates. Statistical test: two-tailed unpaired t-test.

### qRT-PCR

Overnight cultures of *Eco* were washed and OD600-corrected to 1.0 in LB or LBNS +/-100µl IPTG. Three, 10µl aliquots of each culture were applied to nitrocellulose membranes (0.2µm, GVS) atop LB-agar+100µM IPTG or LBNS-agar+100µM IPTG and dried down by flame-sterilization. After growing 6 hours at 30°C, the spots were scalpel-excised, pooled, and resuspended into PBS by bead beating, then pelleted for RNA extraction. RNA was extracted using TRIzol Reagent (Invitrogen) with Max Bacterial Enhancement Reagent (Invitrogen), followed by treatment with Turbo DNA-free kit (Invitrogen) to remove contaminating DNA. After quantification by Nanodrop (Thermo Scientific), total RNA was reverse transcribed into cDNA using qScript cDNA Supermix (QuantaBio). A 1:5 dilution of cDNA and custom primers were input into qPCR reactions with PowerUP SYBR Green Master Mix (Applied Biosystems).qRT-PCR was performed using a QuantStudio 3 Real Time PCR system (ThermoFisher Scientific) using cycling parameters as defined by the master mix instructions. Fold-change in transcript levels was calculated using Λ1Λ1Ct analysis, using *rpoD* as a control gene. Three biological and three technical replicates were used per experiment. Statistical test: two-tailed unpaired t-test. Custom primers for qPCR of *Eco* genes can be found in Table 3.

### Cryo-ET imaging

Overnight cultures of *E.coli* strains were diluted in LB 1:100 and grown at 37°C. At OD600=0.2, 150 µM IPTG was added to the liquid culture to induce CRISPRi knockdown. Bacteria were grown for another 90 min and then flash-frozen in liquid nitrogen. Cell cultures were mixed with 10 nm protein A gold at 20:1 ratio (Utrecht), then aliquots of 3 μL mixtures were applied to glow-discharged R2/2, 200 mesh copper Quantifoil grids (Quantifoil Micro Tools). The sample was blotted for 3 s at 20°C and at 80% humidity. The grids were plunge-frozen in liquid ethane using Leica EM GP system (Leica Microsystems) and stored in liquid nitrogen. Cryo-ET was performed on a Talos electron microscope equipped with a Ceta CCD camera (ThermoFisher). Images were taken at magnification 22,000x corresponding to a pixel size of 6.7 Å. Tilt series were collected using SerialEM^79^ with a continuous tilt scheme (–48° to 48°, every 3° increment). The defocus was set to -6 to -8 μm and the cumulative exposure per tilt series was 150 e^−^/A^2^. Tomograms were reconstructed with the IMOD software package^80^.

### Overexpression plasmid construction and use

Plasmids for periplasmic Tae1 overexpression in *Eco* were constructed using splice-overlap extension cloning of *tae1^WT^* and *tae1^C30A^* coding sequences derived from *P.aeruginosa* (PAO1) into *pBAD24*^48, 81^. A *pelB* leader sequence was fused to *tae1* for localization to the periplasm. Expression from *pBAD24* plasmids transformed into *Eco* was induced by addition of 0.125% arabinose (w/v) (Spectrum Chemical) into liquid LBNS media at early log phase (OD600 ∼0.25). Overexpression constructs for *msbA* and *lpxK* were constructed by cloning each full-length gene from *Eco* into the NdeI/HindIII restriction sites of *pSCrhaB2*^82^. Overexpression from *pSCrhaB2* plasmids transformed into *Eco* was induced by addition of 0.1% rhamnose (w/v) (Thermo Scientific) into liquid media. See **Table S2** for primer sequences used for cloning and PCR validation.

### Tae1 overexpression lysis assay

Chemically competent *Eco* were transformed with Tae1 overexpression constructs (*pBAD24::tae1^WT^, pBAD24::tae1^C30A^, pBAD24*) by standard 42°C heat-shock and a 45-minute recovery in LB at 37°C with shaking. A transformant population was selected overnight in liquid LB+Carb; the more-traditional method of selecting on solid media was skipped to discourage the formation of Tae1-resistant compensatory mutations. Overnight transformant cultures were backdiluted to OD600=0.1 in LBNS+Carb +/-100µM IPTG, then incubated in a Synergy H1 plate reader (BioTek) at 37°C with shaking (2 technical × 3 biological replicates). OD600 reads were taken every five minutes to generate a growth curve. At OD600=0.25 (early log-phase), Tae1 expression was induced from *pBAD24* with the addition of 0.125% arabinose to each well, and grown for 500 minutes at 37°C with shaking. Bacterial growth curves were normalized to blank growth curves (LBNS+Carb, no bacteria), and average growth curves from all biological and technical replicates were plotted in Prism (GraphPad).

For *msbA* and *lpxK* complementation assays*, pSCrhaB2* plasmids were transformed alongside *pBAD24* plasmids, and overnight selection was performed in liquid LB+Carb+Trm. The next day, cultures were washed and backdiluted at OD600=0.1 into LBNS+Carb+Trm+0.1% rhamnose. The experiment then proceeded in the plate reader as described above.

### Western blotting

*dCas9 detection:* Total protein was extracted from the organic layer of bacterial pellets treated with TRIzol Reagent (prepared as described in **qRT-PCR**), according to manufacturer’s protocol. Protein samples were diluted to 1mg/ml in PBS + 1x Laemmli denaturing buffer, boiled for 10 minutes then centrifuged at 20,000xg at RT for 2 minutes. Fifteen µl of supernatant was loaded onto an anyKD MiniPROTEAN gel (BioRad), alongside ProteinPlus Ladder (BioRad). Gels were run according to manufacturer’s protocol in 1x SDS-PAGE running buffer to separate proteins. Protein was transferred to nitrocellulose (0.2µm; GVS) via semi-dry transfer with a TransBlot Turbo transfer system (BioRad) and matching transfer buffer (BioRad) under the following conditions: 45 min @ 15V, 2.5 Amp. Transfer was validated by Ponceau stain. Blots were blocked for one hour at RT with shaking in 3% milk+TBST. Primary antibody was applied: 1:1000 mouse anti-Cas9 (Abcam ab191468) in TBST, overnight, at 4C with shaking. Blots were washed four times in TBST. Secondary antibody was applied: 1:5000 anti-mouse HRP (Advansta R-05071-500) in TBST, for one hour at RT, with shaking. Blots were washed four times in TBST. Blots were treated with Clarity ECL Western blotting substrate (BioRad) for chemiluminescent detection on an Azure c400 imager. Visible light images were also taken to visualize protein ladder. Densitometry analysis was performed in Fiji^83, 84^. Statistical test: two-tailed unpaired t-test. Three biological replicates. *Tae1 detection:* Chemically competent *Eco* cells were transformed with Tae1 overexpression constructs (*pBAD24::tae1^WT^, pBAD24::tae1^C30A^, pBAD24*) by standard 42°C heat-shock and a 45-minute recovery in LB at 37°C with shaking. A transformant population was selected overnight in liquid LB+Carb. Cultures were backdiluted to OD600=0.1 in LBNS + Carb +100µM IPTG, then incubated in a Synergy H1 plate reader (BioTek) at 37°C with shaking (2 technical × 3 biological replicates). OD600 reads were taken every five minutes to track population growth. At OD600=0.25, Tae1 expression was induced with the addition of 0.125% arabinose to each well. Bacteria were grown for 60 minutes with Tae1 induction, before technical replicates were harvested and pooled. Samples were pelleted by centrifugation and media was decanted before cells were resuspended in PBS + 1x Laemmli denaturing buffer. Western blotting protocol then proceeded as above, excepting the use of a custom rabbit anti-Tae1 primary antibody (1:2500 in TBST) (ThermoFisher) and anti-rabbit HRP secondary antibody (1:5000 in TBST) (Advansta R-05072-500).

### Muropeptide analysis

Chemically competent *Eco* cells were transformed with Tae1 overexpression constructs (*pBAD24::tae1^WT^, pBAD24::tae1^C30A^, pBAD24*) by standard 42°C heat-shock and a 45-minute recovery in LB at 37°C with shaking. A transformant population was selected overnight in liquid LB+Carb. Cultures were backdiluted to OD600=0.1 in LBNS+Carb +100µM IPTG, and grown with shaking. At early log phase (OD600=0.25), 0.125% arabinose was added to induce *pBAD24* expression. Cells were grown for 60 minutes, then harvested by centrifugation. For PG purification, cells were boiled in 3% SDS to extract crude PG, then treated with Pronase E (100µg/ml in Tris-HCl (pH 7.2) + 0.06% NaCl) (VWR Chemicals) for 2 hours at 60C to remove proteins covalently bound to PG. Mutanolysin digestion (40µg/ml in Tris-HCl (pH 7.2) + 0.06% NaCl) was performed overnight at 37C to solubilize PG into muropeptides for HPLC analysis. Samples were reduced with sodium borohydride (Fisher Chemical) then pH-corrected to 3-4 using o-phosphoric acid(Fisher Chemical)^85^. Muropeptides were separated on a 1220 Infinity II HPLC (Agilent) with UV-visible detection (λ=206nm). Muropeptide separation was achieved over 54 minutes at 0.5 ml/min using a Hypersil ODS C18 column (Thermo Scientific) and a gradient elution from 50mM sodium phosphate + 0.04% NaN3 (Buffer A) to 75mM sodium phosphate +15% methanol (Buffer B). Chromatograms were integrated in ChemStation software (Agilent) to determine peak area, height, and elution time. Experimental chromatograms were normalized against a chromatogram from a blank run (ddH2O). Chromatograms were also internally normalized against the most abundant M4 (monomer muropeptide) peak; this allowed for direct relative comparisons of peak heights between samples.

To calculate the percent change in D44 (4,3-crosslinked dimer) peptides after Tae1 overexpression, the normalized area under the curve (AUC) for D44 was divided by the total chromatogram area to calculate the relative D44 peak area for each condition (AUCWT, AUCC30A, AUCEV). Then, within a given strain, (AUCWT/AUCEV)*100 and (AUCC30A/AUCEV)*100 were calculated to determine the percent of D44 peak area lost to Tae1^WT^ or Tae1^C30A^ treatment, relative to EV treatment. Three biological replicates were performed per condition. Statistical test: two-tailed unpaired t-test.

### HADA incorporation imaging

Chemically competent cells were transformed with *pBAD24* constructs: (*pBAD24::tae1^WT^, pBAD24::tae1^C30A^*,or *pBAD24*) and selected with Carb overnight in liquid LB. Transformant cultures were backdiluted to OD600=0.1 in 1ml LBNS+Carb +100µM IPTG, and grown with shaking. At early log phase (OD600=0.25), 0.125% arabinose added to induce *pBAD24* expression. Cells were grown for 30 minutes, then 250µM HADA added to culture. Cells were grown an additional 30 minutes, then collected by centrifugation and washed 3x with cold PBS + sodium citrate (pH 3.0) to block hydrolysis of labelled septal PG^86^. Cells were fixed by treatment with 3% PFA for 15 minutes on ice. Fixed cells were washed 3x in cold PBS, then resuspended in PBS +20% DMSO. Fluorescence imaging was performed on a Nikon Eclipse Ti2-E inverted microscope equipped with a 100x/1.40 oil-immersion phase objective and an EMCCD camera (Prime 95B). Fluorescence (DAPI channel) and phase-contrast images were captured using NIS-Elements AR Viewer 5.20. Images were analyzed for single-cell fluorescence intensity using MicrobeJ for Fiji^84, 87^. 200 cells/sample measured, 3 biological replicates. Statistical test: unpaired t-test.

### Nascent protein synthesis imaging

Chemically competent cells were transformed with *pBAD24* constructs: (*pBAD24::tae1^WT^, pBAD24::tae1^C30A^*,or *pBAD24*) and selected with Carb overnight in liquid LB. Cultures were diluted by 1:100 and grown in LBNS+ Carb+ 100µM IPTG at 37 °C with shaking. At early log phase (∼80 minutes) 0.125% arabinose was added to induce Tae1 expression. After 35 minutes, 13µM O-propargyl-puromycin (OPP) was added to cultures to label new peptide synthesis before harvesting (Click-iT™ Plus OPP Alexa Fluor™ 488 Protein Synthesis Assay Kit, Invitrogen)^88^. After labelling, cells were pelleted and fixed in 3.7% formaldehyde in PBS. Cells were permeabilized with 0.3% Triton X-100 in PBS for 15 min, then labelled for imaging with Click-iT reaction cocktail for 20 min in the dark, washed then resuspended in PBS. Fluorescence imaging was performed on a Nikon Eclipse Ti2-E inverted microscope equipped with a 100x/1.40 oil-immersion objective and an EMCCD camera (Prime 95B). The 488-nm laser illumination fluorescence and phase-contrast images were captured using NIS-Elements AR Viewer 5.20 and analyzed using MicrobeJ software for Fiji^84, 87^.

### Time-lapse imaging of T6SS competitions

Competition microscopy experiments were performed with overnight liquid cultures of *Pae* (LB) and *Eco* CRISPRi strains (LB+Gent+Cam). Cultures were diluted 1:50 in fresh medium and grown for 2h. *Pae* cells were diluted again 1:50 in fresh medium (LB) and grown at 37°C to OD 1.2 – 1.5 (∼1 hour). Similarly, *E. coli* strains were diluted 1:100 in fresh medium (LB+150µM IPTG) supplemented with antibiotics (Gent / Cam) and grown at 37°C to OD 1.2 – 1.5 (∼1 hour). Then, cultures were washed with LB, resuspended in LB + 150µM IPTG and mixed 2:1 (*Pae:Eco*). 1 µl of the mixed cells was spotted on an agarose pad containing propidium iodide and imaged for 2h at 37°C. A Nikon Ti-E inverted motorized microscope with Perfect Focus System and Plan Apo 1003 Oil Ph3 DM (NA 1.4) objective lens was used to acquired images. If not indicated otherwise, time-lapse series of competitions were acquired at 10 s acquisition frame rate during 120 min. SPECTRA X light engine (Lumencore), ET-GFP (Chroma #49002) and ET-mCherry (Chroma #49008) filter sets were used to excite and filter fluorescence. VisiView software (Visitron Systems, Germany) was used to record images with a sCMOS camera pco.edge 4.2 (PCO, Germany) (pixel size 65 nm). The power output of the SPECTRA X light engine was set to 20% for all excitation wavelengths. GFP, phase-contrast and RFP / propidium iodide (PI) images were acquired with 50-100 ms exposure time. Temperature and humidity were set to 37°C, 95% respectively, using an Okolab T-unit objective heating collar as well as a climate chamber (Okolab). Fiji was used for imaging processing^84^. Acquired time-lapse series were drift-corrected using a custom StackReg based software ^89, 90^.

## SUPPLEMENTAL INFORMATION

**Table S1**. Bacterial strains and plasmids used in this study

**Table S2**. Primer sequences

**Table S3**. Corrected L2FC values from screen

**Table S4**. Final Diff values from screen

## ACKNOWLEDGEMENTS

We are grateful to all members of the Chou and Basler labs for their support throughout this project (with special thanks to Atanas Radkov, Krisna Van Dyke, Sebastian Flores, and Eleanor Wang). We thank Carol Gross (UCSF) and members of her lab (Jason Peters, Marco Jost, John Hawkins) for their assistance in adapting their CRISPRi system for our project. We thank Michelle Tan, Rene Sit, and Norma Neff (Chan-Zuckerberg Biohub) for assistance with high-throughput sequencing. We thank Naomi Ziv (UCSF), and DeLaine Larsen and Kari Harrington (UCSF Nikon Imaging Center) for assistance with fluorescence microscopy. We are grateful to KC Huang (Stanford University), Waldemar Vollmer (Newcastle University), and Alessandra Polissi (University of Milan) for fruitful conversations regarding the complex biology of the bacterial cell envelope. We thank Sandra Catania and Lauren Trotta for their generous feedback toward data analysis and manuscript preparation.

This work was funded by: NIH NIAID award T32AI060535 (KLT), a UCSF Moritz-Heyman Discovery Fellowship (KLT), the Swiss National Science Foundation (grant BSSGI0_155778) (MB), a European Research Council consolidator grant (865105 -“AimingT6SS”) (MB), the Chan-Zuckerberg Biohub (SC), and the Pew Biomedical Scholars Program (SC).

## AUTHOR CONTRIBUTIONS

Kristine L Trotta (Conceptualization, Methodology, Research, Data analysis, Data visualization, Writing, Reviewing)

Beth M Hayes (Conceptualization, Research, Data analysis, Reviewing)

Johannes P Schneider (Methodology, Research, Data analysis, Reviewing)

Jing Wang (Methodology, Research, Data analysis, Reviewing)

Horia Todor (Data analysis, Data visualization, Reviewing)

Patrick Rockefeller Grimes (Research, Reviewing)

Ziyi Zhao (Methodology, Research, Data analysis, Reviewing)

William L Hatleberg (Data visualization, Reviewing)

Melanie R Silvis (Conceptualization, Methodology, Reviewing)

Rachel Kim (Methodology, Reviewing)

Byoung-Mo Koo (Methodology, Reviewing)

Marek Basler (Project administration, Reviewing)

Seemay Chou (Project administration, Funding, Conceptualization, Writing, Reviewing)

## SUPPLEMENTAL FIGURES

**Supplement 1:**
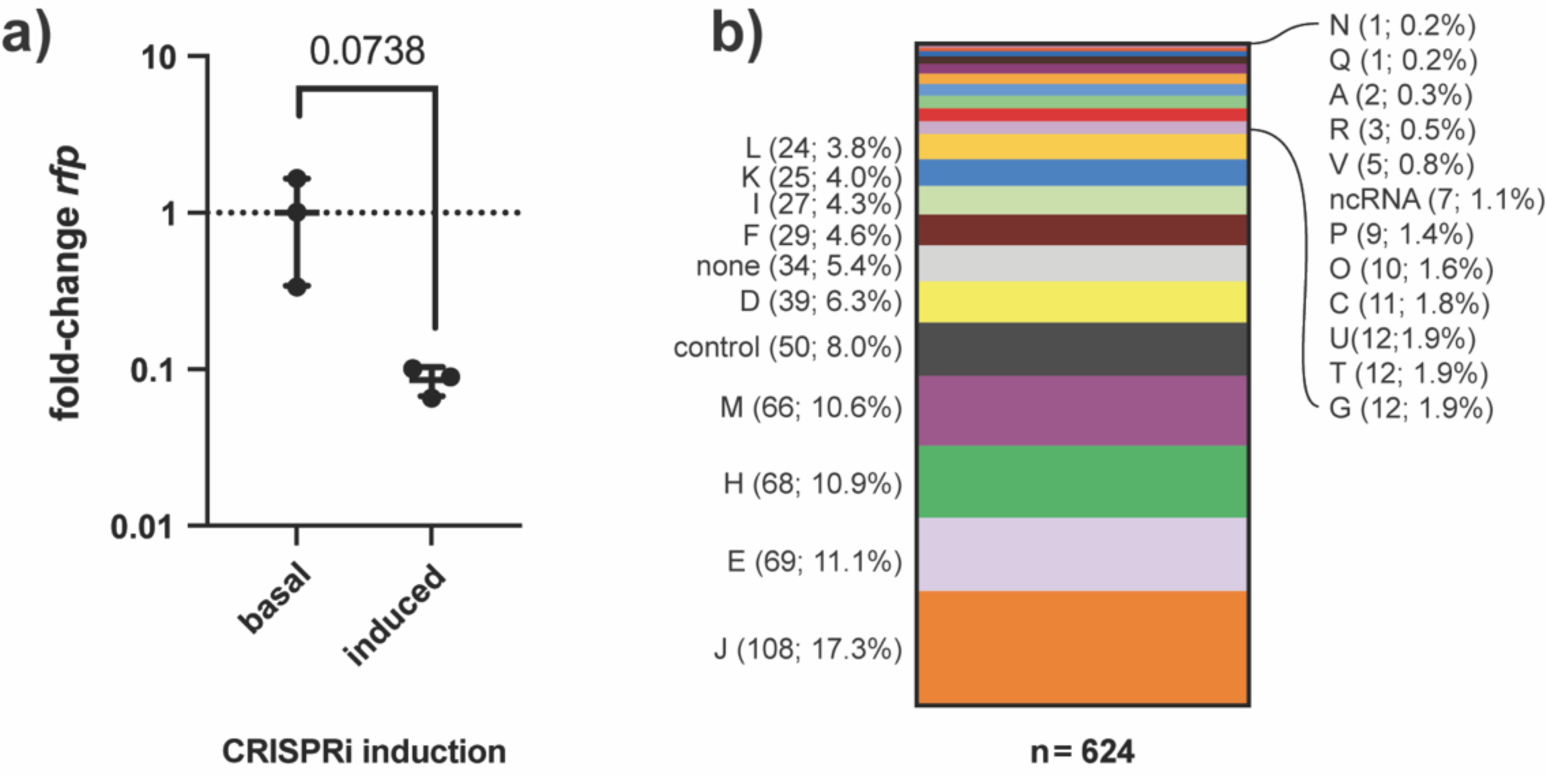
CRISPRi conditionally knocks down transcription across hundreds of *Eco* gene targets. **a) CRISPRi induction produces mild transcriptional knockdown of endogenous *rfp* (11.7-fold decrease) in *Eco***. qRT-PCR measurement of relative *rfp* RNA expression in *Eco* strain SC363 after 6 hours of growth on solid LB media with basal or induced CRISPRi. Data shown: 3 biological replicates with mean ± s.d. Statistical test: unpaired two-tailed *t*-test. **b) CRISPRi targets *Eco* genes that collectively represent 21 clusters of orthogonal genes (COGs).** CRISPRi target genes (*n*=596) were binned by their NCBI COG functional assignment. The relative representation of each COG in the strain collection is displayed as a percent of all COGs. Some genes are represented by multiple COGs, resulting in a greater number of COGs (*n*=624) than target genes. Non-targeting negative controls (“control”, *n*=50) genes without COG assignments (“none”, *n*=34), and genes coding for non-coding RNAs (“ncRNA”, *n*=7) were also binned.

**Supplement 2:**
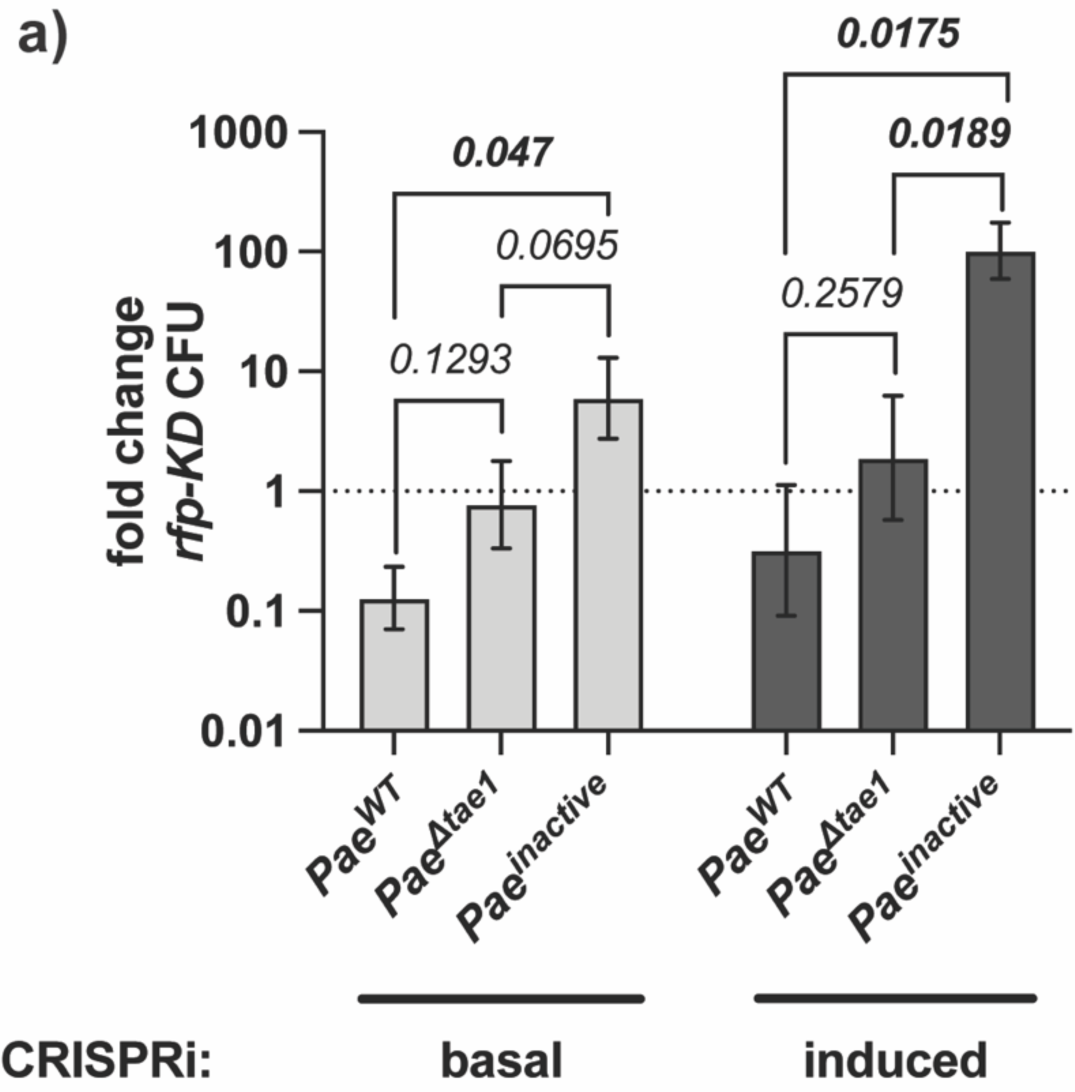
Non-targeting CRISPRi induction has little effect on *Eco* fitness in T6SS competition. **a) CRISPRi induction does not disrupt T6SS- and Tae1-dependent targeting of *Eco* by *Pae*.** Interbacterial competition between *Pae* (*Pae^WT^, Pae ^Δtae^*^1^*, Pae^inactive^*) and an *Eco* negative-control KD strain (*rfp-KD*), with induced or basal CRISPRi. Data shown: mean fold-change (± geometric s.d.) of *rfp-KD* colony forming units (CFUs) after six hours of competition against *Pae*. Statistical test: unpaired two-tailed *t-*test; *p*-value ≤0.05 displayed in bold font.

**Supplement 3:**
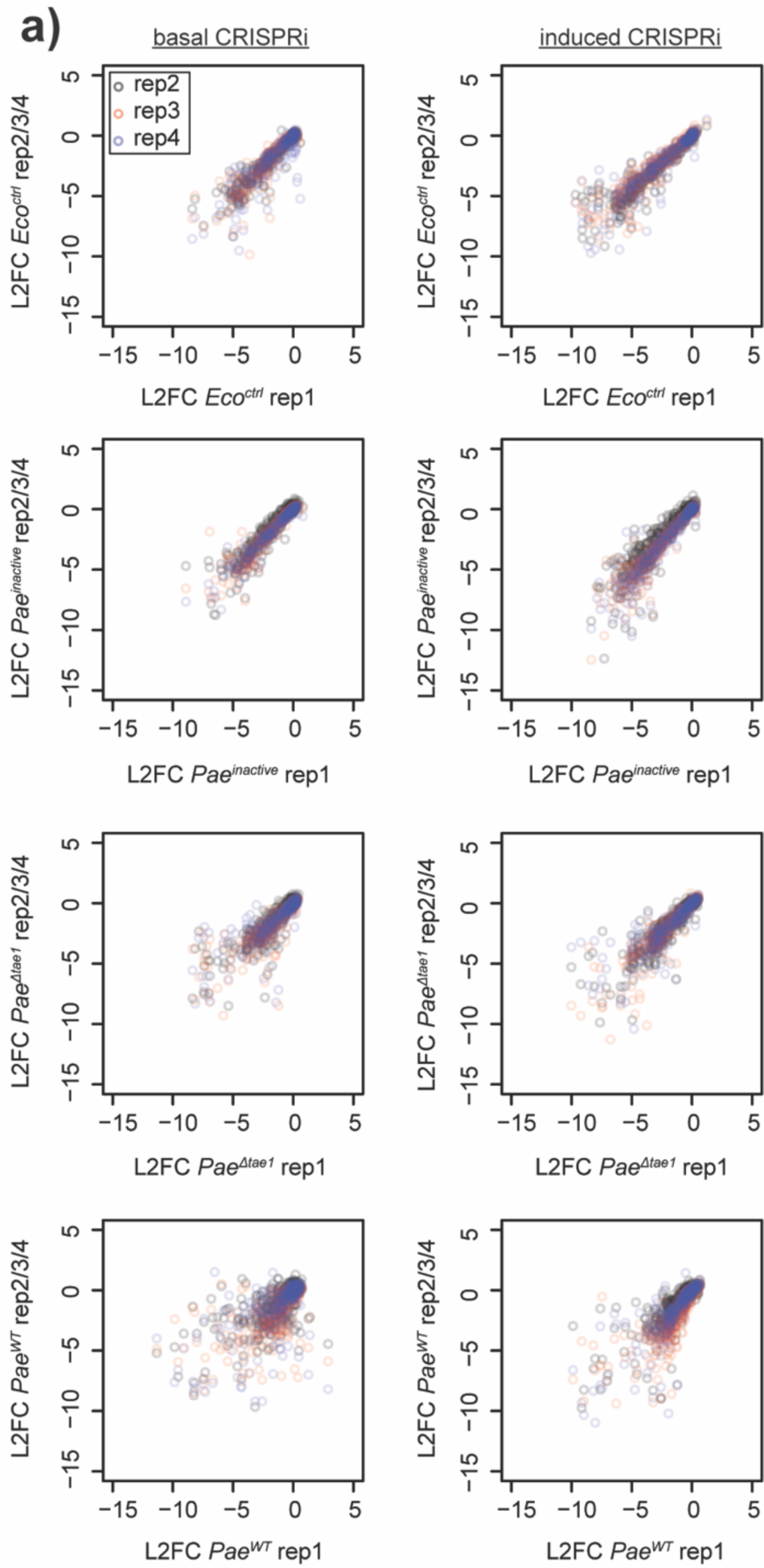
CRISPRi library fitness in T6SS screen is reproducible across biological replicates. **a) CRISPRi library fitness in T6SS screen is reproducible across biological replicates.** Replica plots showing the uncorrected L2FC values for each *Eco* CRISPRi strain after competition against *Pae^WT^, Pae ^Δtae^*^1^*, Pae^inactive^*, for four biological replicates. For each plot, replicate 1 is compared to replicate 2 (grey), replicate 3 (red), or replicate 4 (blue). Median Pearson’s *r* between all replicates = 0.91.

**Supplement 4:**
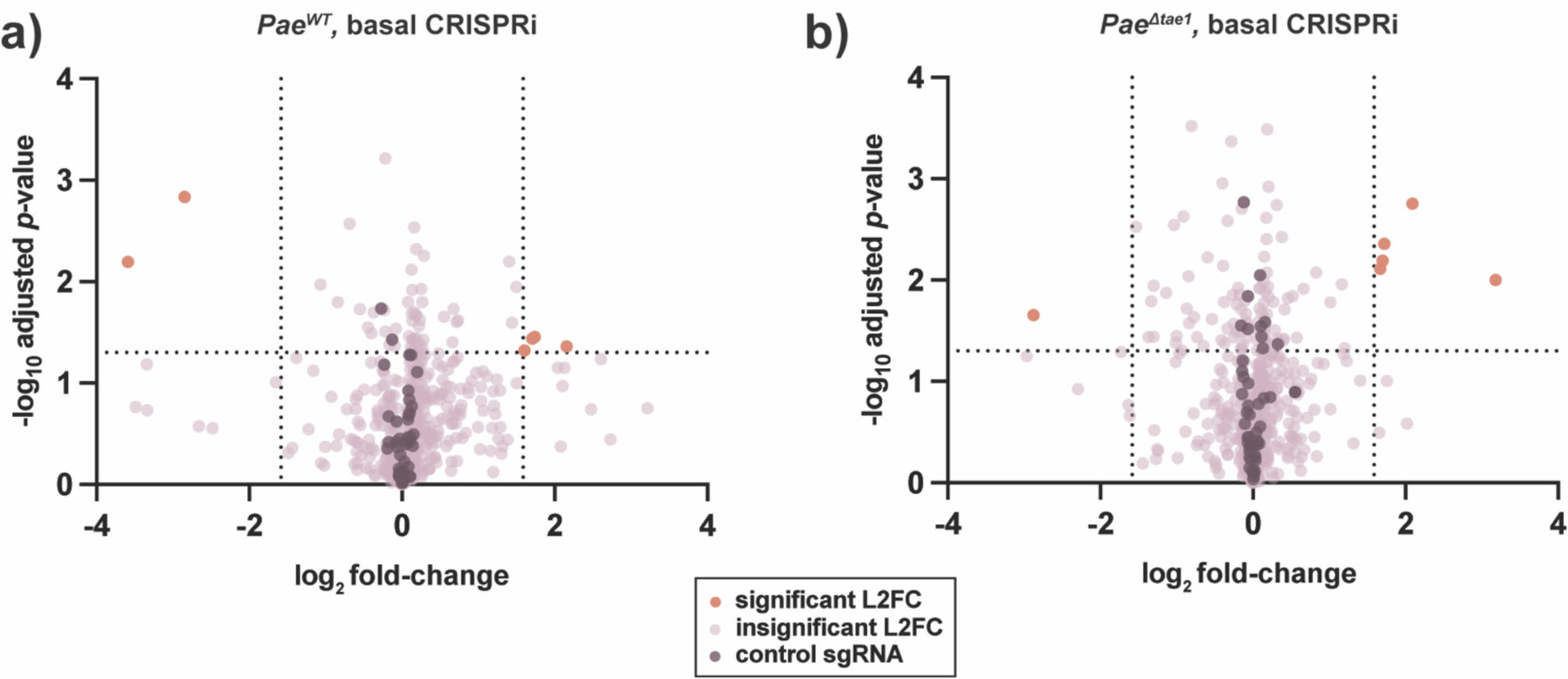
Pooled T6SS competitions with basal CRISPRi attenuate significant fitness phenotypes. **a-b) Basal CRISPRi attenuates *Eco* fitness phenotypes against *Pae^WT^* (a) and *Pae^Δtae^*^1^(b).** Volcano plots showing log2-fold change (L2FC) values for each KD strain after interbacterial competition (basal CRISPRi). Data shown: mean from four biological replicates. Statistical test: Wald test. Vertical dotted lines indicate arbitrary cutoffs for L2FC at x =-1.58 and x=1.58 (absolute FC x=-3 or x= 3). Horizontal dotted line indicates statistical significance cutoff for log10 adjusted p-value (≤ 0.05). Orange points represent KDs with L2FC ≥ 1.58 or ≤ -1.58 and log10-adj. *p*-value ≤0.05. Dark purple points represent non-targeting negative control KDs (*n*=50). Lavender points represent KDs that do not meet cutoffs for L2FC or statistical test.

**Supplement 5:**
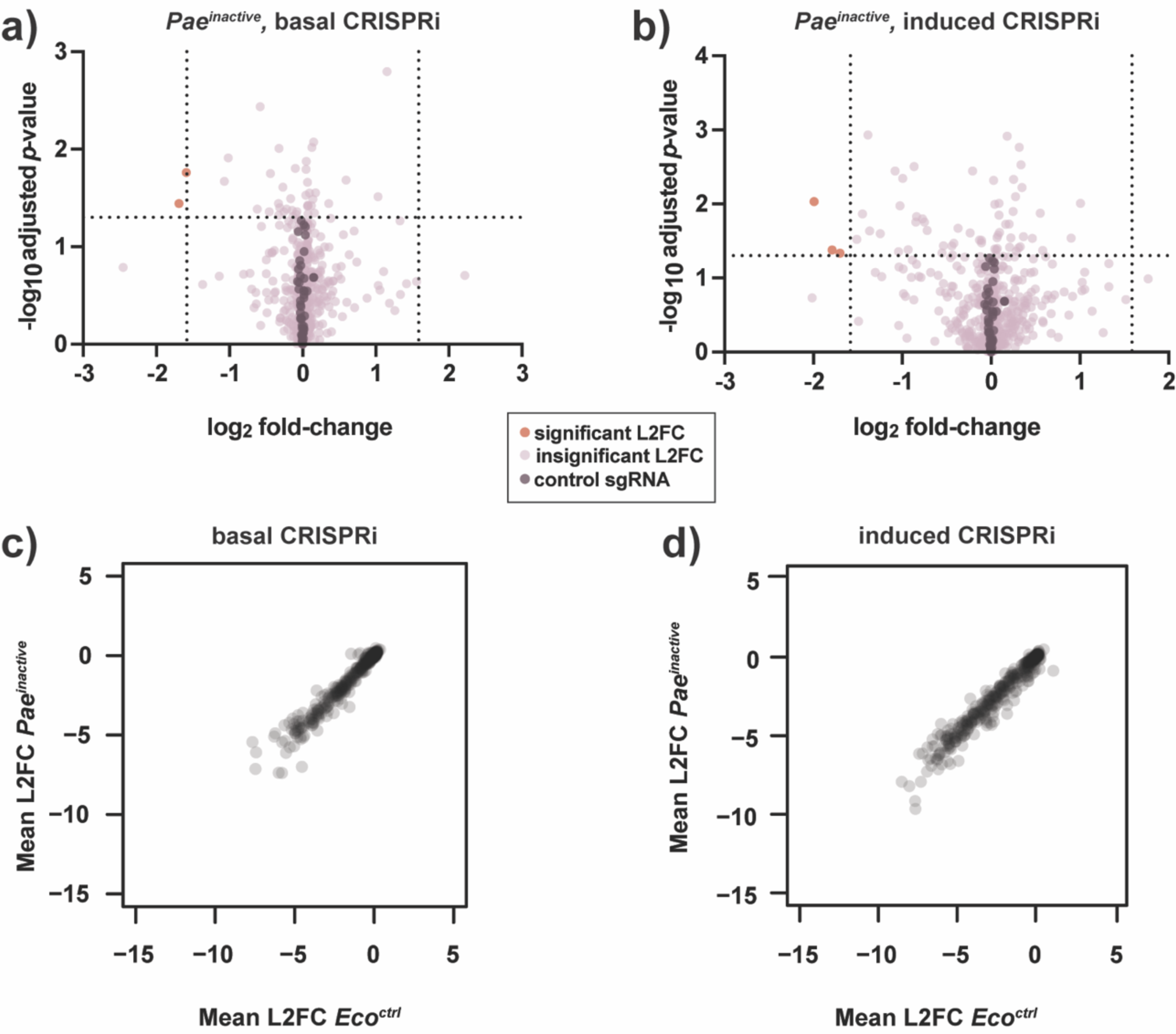
*Pae^inactive^* is a neutral co-culture partner for *Eco*. **a-b) Competition against *Pae^inactive^* reveals few *Eco* fitness determinants.** Volcano plots showing log2-fold change (L2FC) values for each KD strain after interbacterial competition with induced (a) or basal (b) CRISPRi. Data shown: mean from four biological replicates. Statistical test: Wald test. Vertical dotted lines indicate arbitrary cutoffs for L2FC at x =-1.58 and x=1.58 (absolute FC x=-3 or x= 3). Horizontal dotted line indicates statistical significance cutoff for log10 adjusted p-value (≤ 0.05). Orange points represent KDs with L2FC ≥ 1.58 or ≤ -1.58 and log10-adj. *p*-value ≤0.05. Dark purple points represent non-targeting negative control KDs (*n*=50). Lavender points represent KDs that do not meet cutoffs for L2FC or statistical test. **c-d) KD strain abundance is highly similar after competition with *Pae^inactive^* and after growth without competition (*Eco^ctrl^*).** Scatter plots comparing mean L2FC for each *Eco* KD strain after competition with *Pae^inactive^* or *Eco^ctrl^* treatment, with basal (c) or induced (d) CRISPRi. Median Pearson correlation *r*= 0.98.

**Supplement 6:**
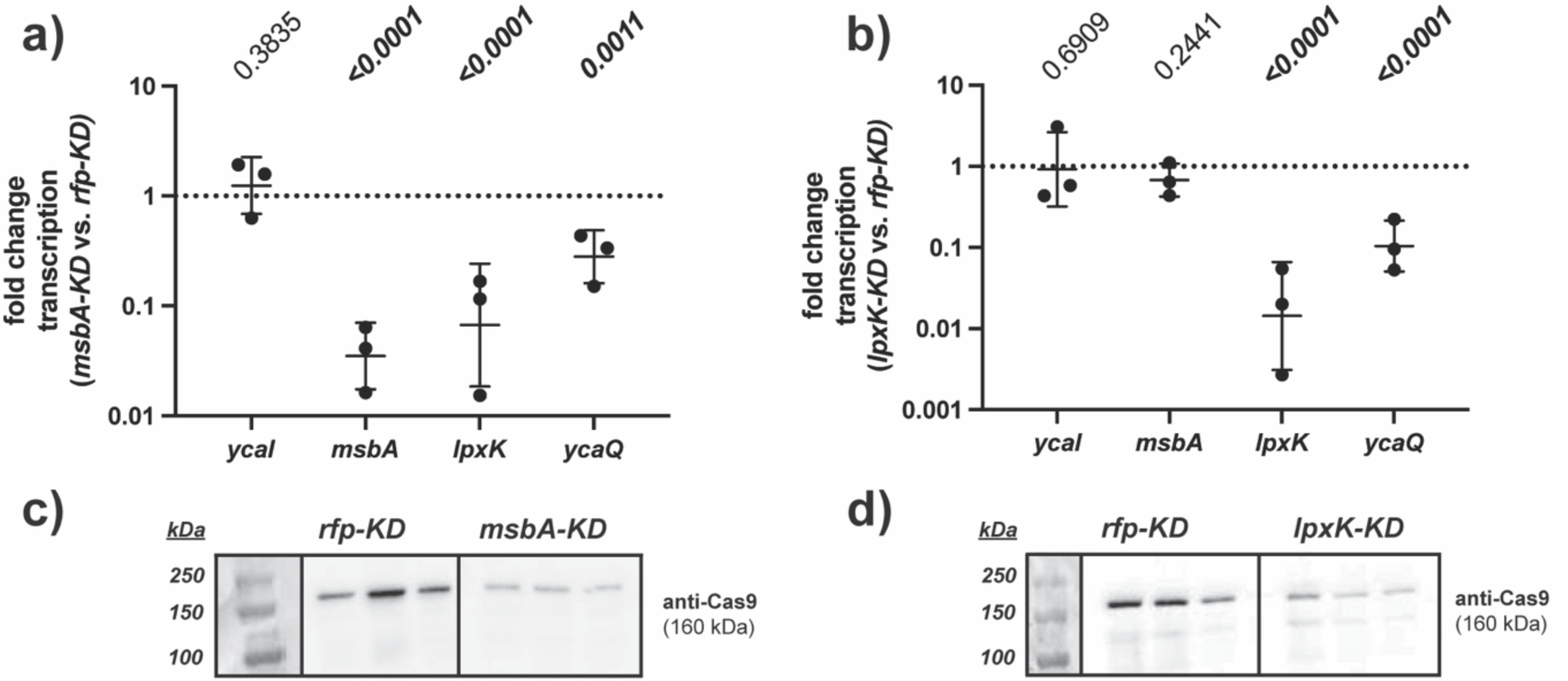
*lpxK-KD* and *msbA-KD* modulate target gene expression and show polar effects. **a-b) Transcriptional knockdowns in *msbA* and *lpxK* have off-target polar effects on transcription in their operon.** qRT-PCR analysis of transcriptional fold-change in *ycaI-msbA-lpxK-ycaQ* in *msbA-KD* (a) and in *lpxK-KD* (b) after growth for 6 hours with induced CRISPRi, normalized to expression in *rfp-KD*. Data shown are geometric average of 3 biological replicates ± s.d. Statistical test: unpaired two-tailed *t*-test; *p*-value ≤0.05 displayed in bold font. **c-d) *msbA-KD* and *lpxK-KD* express a catalytically dead Cas9 (dCas9) enzyme for CRISPRi-mediated transcriptional knockdown.** Western blot analysis of dCas9 protein expression (160 kDa) from *rfp-KD, msbA-KD* (c), and *lpxK-KD* (d). Three independent biological replicates shown.

**Supplement 7:**
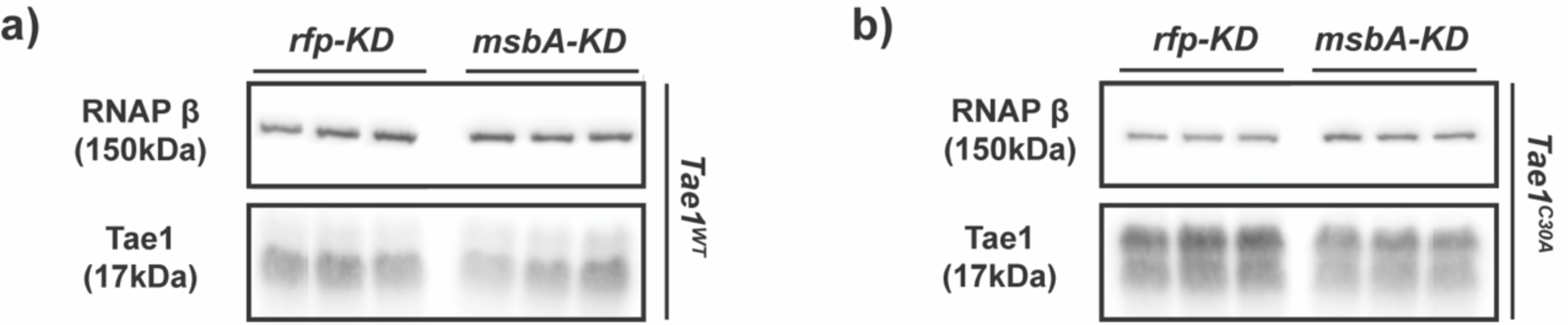
Tae1 protein expression is unaffected in *msbA-KD*. **a-b) Bulk Tae1 protein expression is similar between *msbA-KD* and *rfp-KD*.** Western blot analysis of periplasmic Tae1 protein (17kDa) from (a) *pBAD24::pelB-tae1^WT^*(Tae1^WT^) or (b) *pBAD24::pelB-tae1^C30A^* (Tae1^C30A^) in *rfp-KD* and *msbA-KD* (with induced CRISPRi). Protein expression of RNA polymerase (β subunit) (150kDa) is used as an internal loading control.

**Supplement 8:**
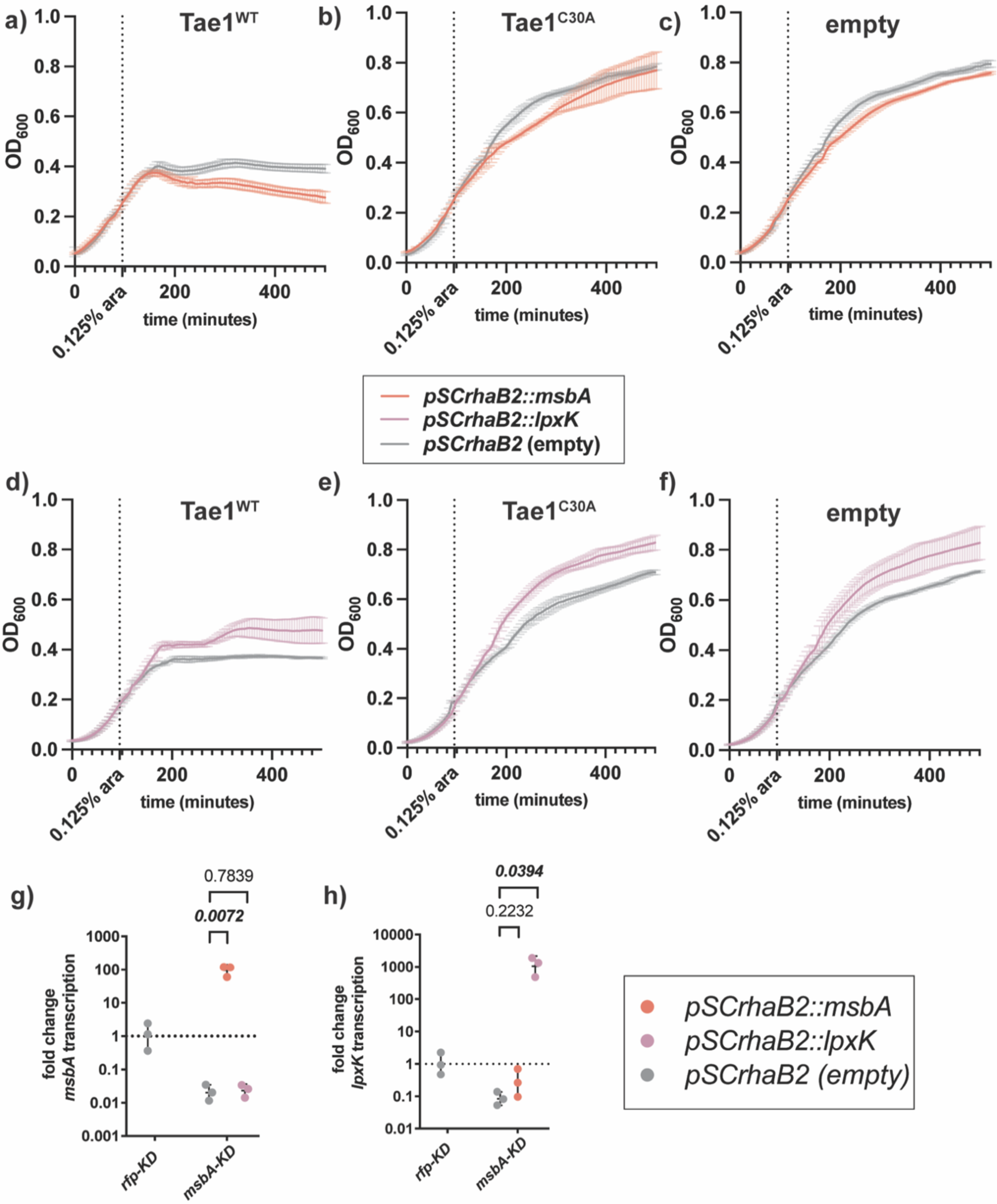
Plasmid-borne overexpression of *msbA* partially rescues Tae1 sensitivity in *msbA-KD*. **a-c) Plasmid-borne *msbA* overexpression partially rescues *msbA-KD* resistance to lysis by Tae1**. OD600 growth curves of *msbA-KD* with induced CRISPRi, overexpressing *pSCrhaB2::msbA* (orange) or *pSCrhaB2* (empty) (grey) alongside (a)*pBAD24::pelB-tae1^WT^* (Tae1^WT^), (b) *pBAD24::pelB-tae1^C30A^* (Tae1^C30A^), or (c) *pBAD24* (empty). Data shown: average of 3 biological replicates ± s.d. Dotted vertical line indicates *pBAD24* induction timepoint (at OD600=0.25) (0.125% arabinose w/v). **d-f) Plasmid-borne *lpxK* overexpression enhances *msbA-KD* resistance to lysis by Tae1.** OD600 growth curves of *msbA-KD* with CRISPRi induced, overexpressing *pSCrhaB2::lpxK* (purple) or *pSCrhaB2* (empty) (grey) alongside (d)*pBAD24::pelB-tae1^WT^* (Tae1^WT^), (e) *pBAD24::pelB-tae1^C30A^*(Tae1^C30A^), or (f) *pBAD24* (empty). Data shown: average of 3 biological replicates ± s.d. Dotted vertical line indicates *pBAD24* induction timepoint (at OD600=0.25) (0.125% arabinose w/v). **g-h) *pSCrhaB2* vectors selectively rescue transcription of their target gene by overexpression.** qRT-PCR analysis of transcriptional fold-change in (g)*msbA* or (h)*lpxK* expression with constitutive rhamnose induction of *pSCrhaB2::msbA* (orange), *pSCrhaB2::lpxK* (purple), or (c)*pSCrhaB2* (empty; grey) in *msbA-KD* with induced CRISPRi. Expression normalized against basal *msbA* expression in *rfp-KD* + *pSCrhaB2* (empty). Data shown: geometric average of 3 biological replicates ± s.d. Statistical test: unpaired two-tailed *t*-test; *p*-value ≤0.05 displayed in bold font.

**Supplement 9:**
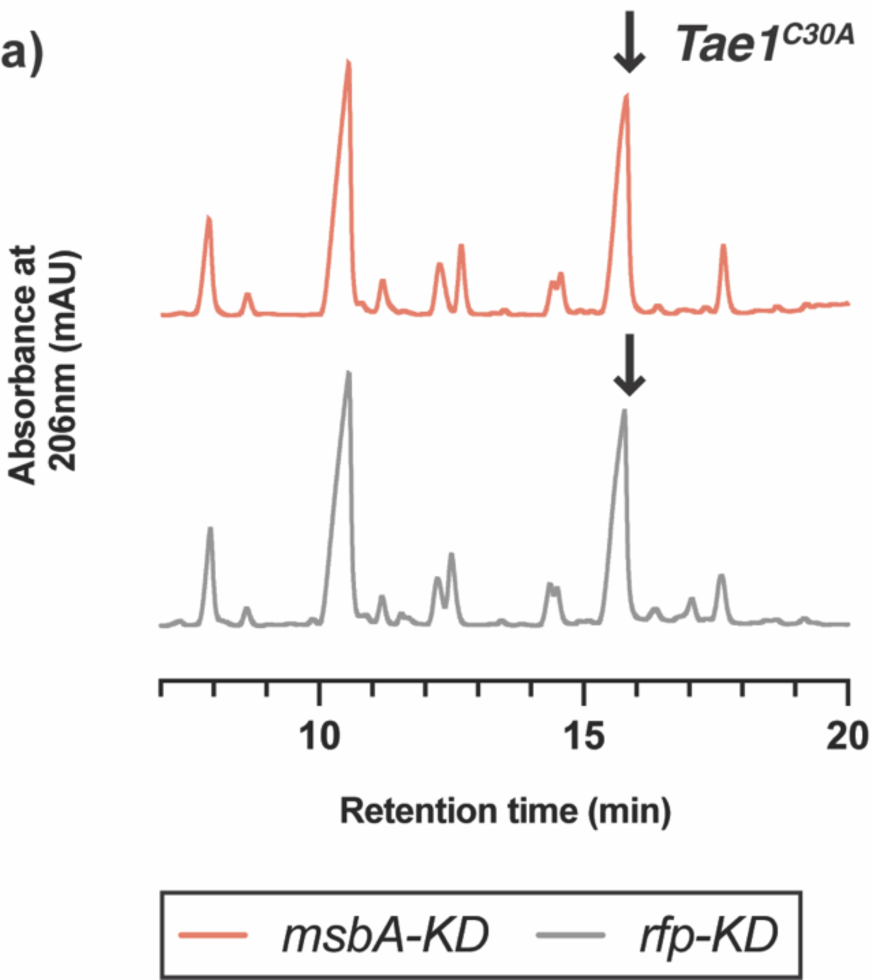
Tae1^C30A^ hydrolyzes D44 muropeptides in *rfp-KD* and *msbA-KD*. **a)Tae1^C30A^ overexpression yields minor digestion of D44 muropeptides**. HPLC chromatograms of muropeptides purified from *msbA-KD* (orange) and *rfp-KD* (grey) expressing *pBAD24::pelB-tae1^C30A^* (Tae1^C30A^). Black arrow indicates D44 peptide partially digested by Tae1^C30A^. Data shown: representative from 3 biological replicates.

**Supplement 10:**
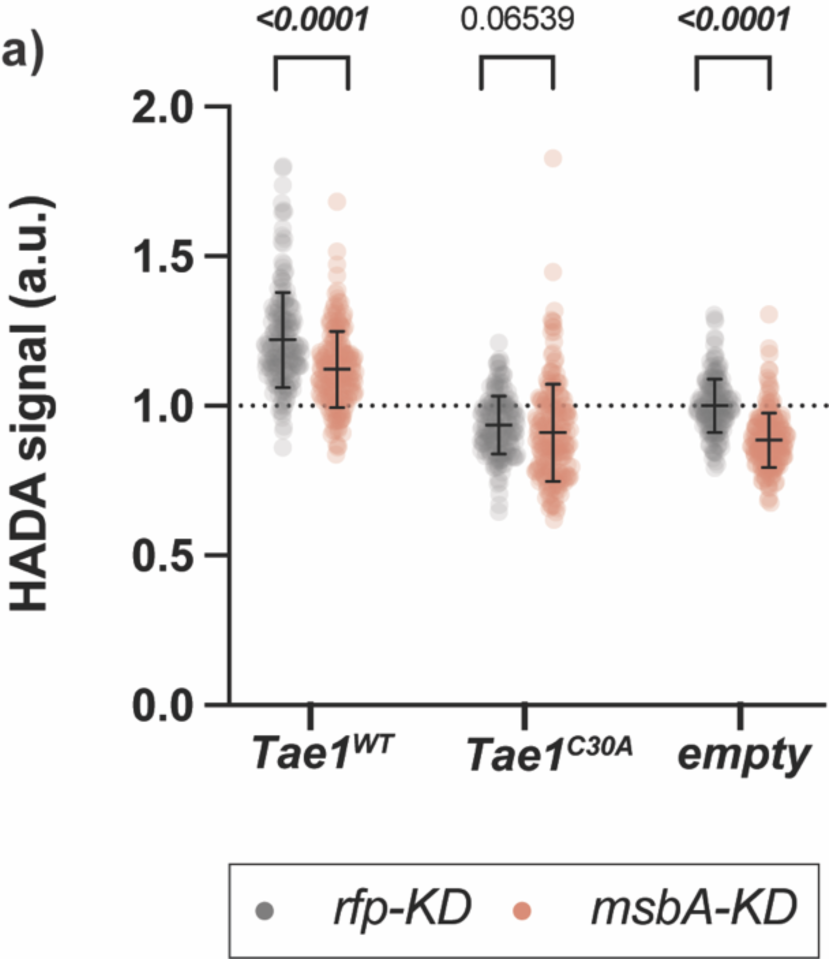
PG synthesis activity is *msbA-KD* is suppressed across all conditions. **a) PG synthesis activity in *msbA-KD* is attenuated under all tested conditions.** Single-cell fluorescence intensity measurements for *rfp-KD* (grey) or *msbA-KD* (orange) incorporating the fluorescent D-amino acid HADA into PG after 60 minutes of overexpressing *pBAD24::pelB-tae1^WT^*(Tae1^WT^), *pBAD24::pelB-tae1^C30A^* (Tae1^C30A^), or *pBAD24* (empty), with CRISPRi induced. All data normalized to average HADA signal in *rfp-KD +* empty. Data shown: 600 cells (200 cells x 3 biological replicates), with average ± s.d. Statistical test: unpaired two-tailed *t*-test; *p*-value ≤0.05 displayed in bold font.

**Supplement 11:**
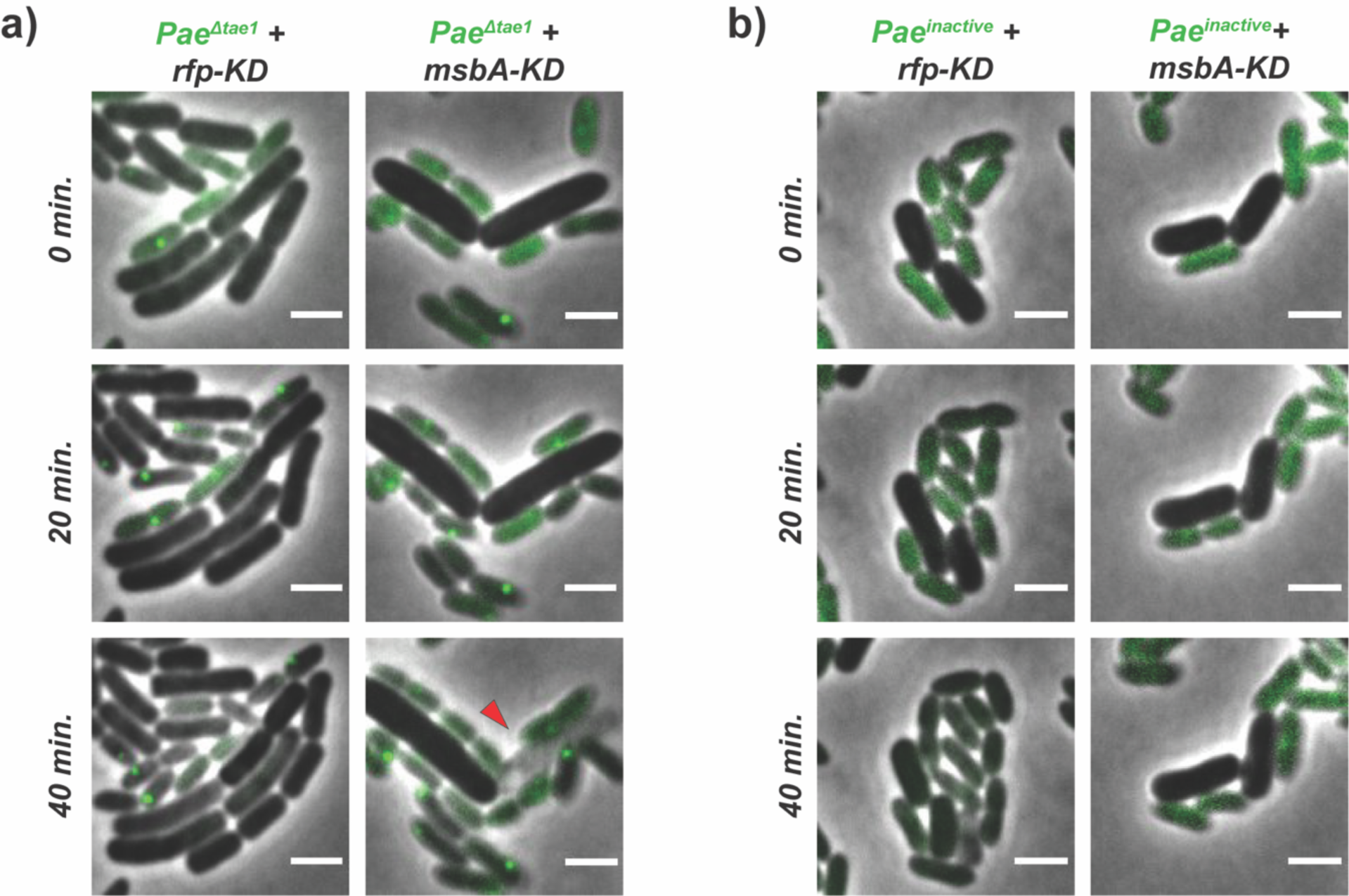
*msbA-KD* growth defects are independent of *Pae* T6SS activity. ***a-b*) *msbA-KD* cells maintain growth defects regardless of *Pae* competitor**. Representative frames from time-course imaging of *rfp-KD* (left column; grey cells) and *msbA-KD* (right column; grey cells) co-cultured with *Pae ^Δtae^*^1^(a) or *Pae ^inactive^* (b) (green cells), and with induced CRISPRi. Green foci in *Pae^WT^* indicate aggregates of GFP-labelled ClpV, which signal H1-T6SS firing events. Red arrow indicates lysed cell. Data shown are merged phase and fluorescent channels. Scale bar: 2µm.

